# Monosynaptically-interconnected Network Module (MNM) Approach for High-Resolution Brain Sub-Network Analysis

**DOI:** 10.1101/2024.02.19.581007

**Authors:** Sunwhi Kim, Yong-Eun Kim, Yusuke Ujihara, Il Hwan Kim

## Abstract

We introduce the Monosynaptically-interconnected Network Module (MNM) approach, an innovative method designed for efficiently analyzing the anatomical structure and functional dynamics of specific brain network modules *in vivo*. Utilizing an Intein-mediated split-Cre system combined with bidirectional adeno-associated viruses, this technique precisely targets and manipulates monosynaptically interconnected modular subnetworks in freely moving animals. We demonstrate its utility through anatomical and functional mapping of a specific MNM encompassing the prefrontal cortex (PFC), basolateral amygdala (BLA), and intermediary hub regions. Specifically, the MNM approach with Cre-reporter mice visualizes detailed network architecture and enables the tracing of axonal connections among the nodes in the network. Furthermore, integration of the MNM approach with Cre-dependent Ca^2+^ indicator and multi-fiber photometry in freely moving mice reveals enhanced correlative network activities in social contexts. This versatile technique offers significant potential for advancing our understanding of network functions that underlie complex behaviors, providing a modular network perspective.

## Introduction

Recent advances in network neuroscience have shifted our research focus from perceiving the brain as a collection of separate regions to understanding it as a complex network of interconnected areas^1–6^. Studies suggest that the brain is composed of smaller subnetworks or modules, densely linked within but loosely between each other, forming a small-world network^7–13^. This modular view serves as a useful framework for understanding how multiple brain regions collaborate effectively, contributing to overall brain activity and behavior. However, current methods in network neuroscience face significant challenges in accurately identifying/visualizing neural network modules. Diffusion tensor imaging (DTI) models the whole-brain connectome by mapping white matter tracts^14, 15^ but is limited in resolution to distinguish individual axonal pathways from complex intersections of fiber tracts^16^, leading to potential misinterpretation of the modules. Functional magnetic resonance imaging (fMRI) and electrophysiological techniques can estimate brain modules through correlated activities^17–26^, yet these methods lack the anatomical specificity to confirm findings^20^. While there have been efforts to combine the structural and functional modules^11, 27^ it has also been reported they are not necessarily aligned^28^, leaving a fundamental gap between the two methods. Recently, tracer-based methods using recombinant adeno-associated viruses (rAAVs) have significantly improved the elucidation of axonal projections^29–35^. Nonetheless, the outcomes of these approaches have frequently been ambiguous due to the extremely complex axonal signals spreading across the brain with afferent or efferent projections. To overcome these challenges, a new and innovative tool is needed – one that can precisely analyze the architecture of specific network modules, monitor their activities, and manipulate genes of interest within the network.

Here we introduce the Monosynaptically-interconnected Neural Network Module (MNM) approach, a novel technique developed from our previously established strategy for selecting single long-range neural circuits^36–39^. The ‘MNM’ concept refines the broader term ‘module’ in network neuroscience, offering more precise identification of monosynaptically-interconnected distant brain regions *via* axons. MNM tracing utilizes two rAAV vectors – anterograde trans-synaptic rAAV2/1^30, 32–34, 40^ and retrograde rAAV2.retro^31^ – each carrying components of the ‘Intein-fused split-Cre’^41^. This design enables convergence of rAAV vectors within neurons that are monosynaptically-interconnected across distant brain regions. This convergence triggers an Intein-mediated reconstitution process, resulting in the formation of the full-length Cre recombinase enzyme within these neurons. Subsequently, this allows for the selective illumination of targeted modular networks in the Cre-reporter mouse model.

In this study, we demonstrate the utility of MNM approach by mapping a specific MNM, termed MNM^PFC→BLA^, that is composed of the prefrontal cortex (PFC), basolateral amygdala (BLA), and intermediary hub regions such as the paraventricular nucleus of the thalamus (PVT) and insular cortex (IC). We achieved detailed visualization of the MNM^PFC→BLA^ topology using whole-brain clearing, light-sheet imaging, and 3-D reconstruction. In addition, we monitored real-time neural activity of the network during social behavior using network-selective Cre-dependent Ca^2+^ indicator (GCaMP8m) transduction and multi-fiber photometry. The results revealed an increase in correlative neural activities in the network during social behavior. These findings affirm the MNM approach as a robust and groundbreaking tool for conducting comprehensive structural and functional analyses of modular networks *in vivo*.

## Results

### Comparison of MNM approach with conventional rAAV tracing methods

We first compared our MNM approach with conventional rAAV tracing methods. rAAV2/1 (often abbreviated as AAV1) is one of the most widely used rAAV serotypes for anterograde trans-synaptic tracing^30, 32–34, 40, 42^. We injected rAAV2/1-hSyn-*Cre* into the right PFC of a Cre-reporter Ai-14 mouse (Fig. 1a). Brain clearing and light-sheet imaging revealed extensive tdTomato fluorescence reporting Cre in brain-wide projections from the right PFC, extending to numerous regions across both hemispheres (Fig. 1b, Extended Fig. 1, Video 1). Notably, due to the trans-synaptic traveling property of rAAV2/1^32–34, 40^, tdTomato signals were also observed in the neuronal cell bodies of these recipient regions. This result demonstrates the effectiveness of conventional anterograde rAAV tracing; however, it also highlights its lack of selectivity, as evidenced by the broad array of labeled projections.

**Figure 1.**
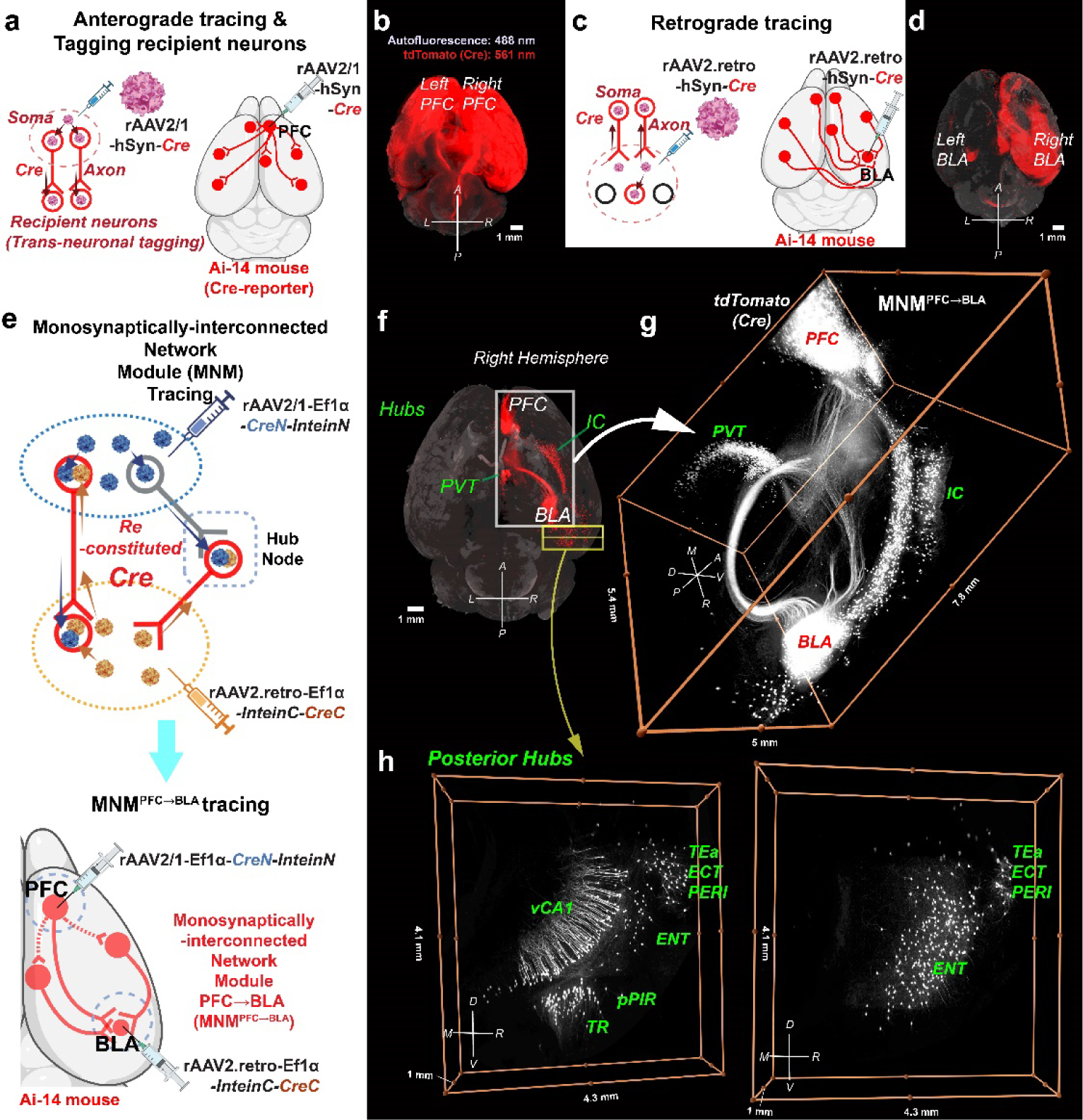
Comparative analysis of conventional recombinant Adeno-Associated Virus (rAAV) and Monosynaptically-interconnected Network Module (MNM) tracing techniques. (a) Schematics of anterograde tracing and tagging of the recipient neurons, using rAAV2/1-hSyn-Cre: (left) rAAV2/1-hSyn-Cre’s properties as an anterograde tracer with trans-synaptic transport capabilities; (right) its application in tracing from the right prefrontal cortex (PFC) and tagging recipient neurons. (b) Cleared brain from an Ai-14 (Cre-reporter with tdTomato) mouse injected with rAAV2/1-hSyn-Cre in the right PFC. The red tdTomato fluorescence, indicating Cre recombinase activity (excited at 561 nm), demonstrates anterograde tracing from the right PFC to its recipient neurons. Background autofluorescence of the brain structure is shown in gray (excited at 488 nm). (c) Schematics of retrograde tracing with rAAV2.retro-hSyn-Cre: (left) rAAV2.retro-hSyn-Cre’s properties as a retrograde tracer in addition to marking neurons at the injection site; (right) its application in tracing afferent projections to the right basolateral amygdala (BLA). (d) Cleared brain from an Ai-14 mouse injected with rAAV2.retro-hSyn-Cre in the right BLA. The tdTomato signals indicate the right BLA and the regions that project to the right BLA. (e) Overview of the (Monosynaptically-interconnected Network Module) MNM Tracing Technique: (Top) Schematic of the MNM tracing approach, which employs a combination of rAAV2/1 and rAAV2.retro vectors with a split-Cre system, enabling selective Cre reconstitution across distant brain regions and intermediary hub regions that comprise a modular network. (Bottom) Application of MNM tracing in the right hemisphere, specifically targeting the MNM centered on the PFC→BLA circuit (MNMPFC→BLA). In Ai-14 mice, rAAV2/1-Ef1α-CreN-InteinN is injected into the right PFC, and rAAV2.retro-Ef1α-InteinC-CreC into the right BLA. The rAAV vectors’ capacity for anterograde and retrograde transport of split-Cre components, coupled with the trans-synaptic travel property of rAAV2/1, ensures Cre reconstitution within neurons of both the direct PFC→BLA circuit and associated hub regions, effectively delineating the MNMPFC→BLA. (f) Cleared brain from an Ai-14 mouse with MNMPFC→BLA tracing in the right hemisphere. The tdTomato signals are observed in the PFC and BLA, along with hub regions including the paraventricular nucleus of the thalamus (PVT), insular cortex (IC), and various regions in the posterior part of the right hemisphere. (g) Detailed view of the MNMPFC→BLA. The tdTomato signals are converted to white to emphasize axonal connections. This MNM tracing effectively delineated the axonal bundles linking the PFC, PVT, IC, and BLA. (h) Detailed view of the hub regions in posterior part of the right hemisphere. These images provide a closer look at the posterior hub nodes of the MNMPFC→BLA in the right hemisphere, specifically highlighting regions such as the temporal association areas (TEa), ectorhinal area (ECT), perirhinal area (PERI), entorhinal area (ENT), posterior piriform area (pPIR), postpiriform transition areas (TR), and ventral hippocampus CA1 (vCA1). Notably, the axonal connections in these posterior hubs are sparser and less distinct than those observed in the anterior part of the network. Note: Directional markers indicate the following orientation: A (anterior), P (posterior), L (left), R (right), M (medial), D (dorsal), and V (ventral).

In a parallel approach, we injected the retrograde tracer rAAV2.retro^29, 31, 43^, carrying the Cre recombinase (rAAV2.retro-hSyn-*Cre*), into the right BLA of an Ai-14 mouse (Fig. 1c). This approach successfully delineated afferent pathways, yet it revealed intricate projections without clear selectivity. This lack of selectivity was apparent in the various illuminated input regions throughout the brain, many of which overlapped with the regions identified through anterograde tracing from the PFC (Fig. 1d, Extended Fig. 2, Video 2).

**Figure 2.**
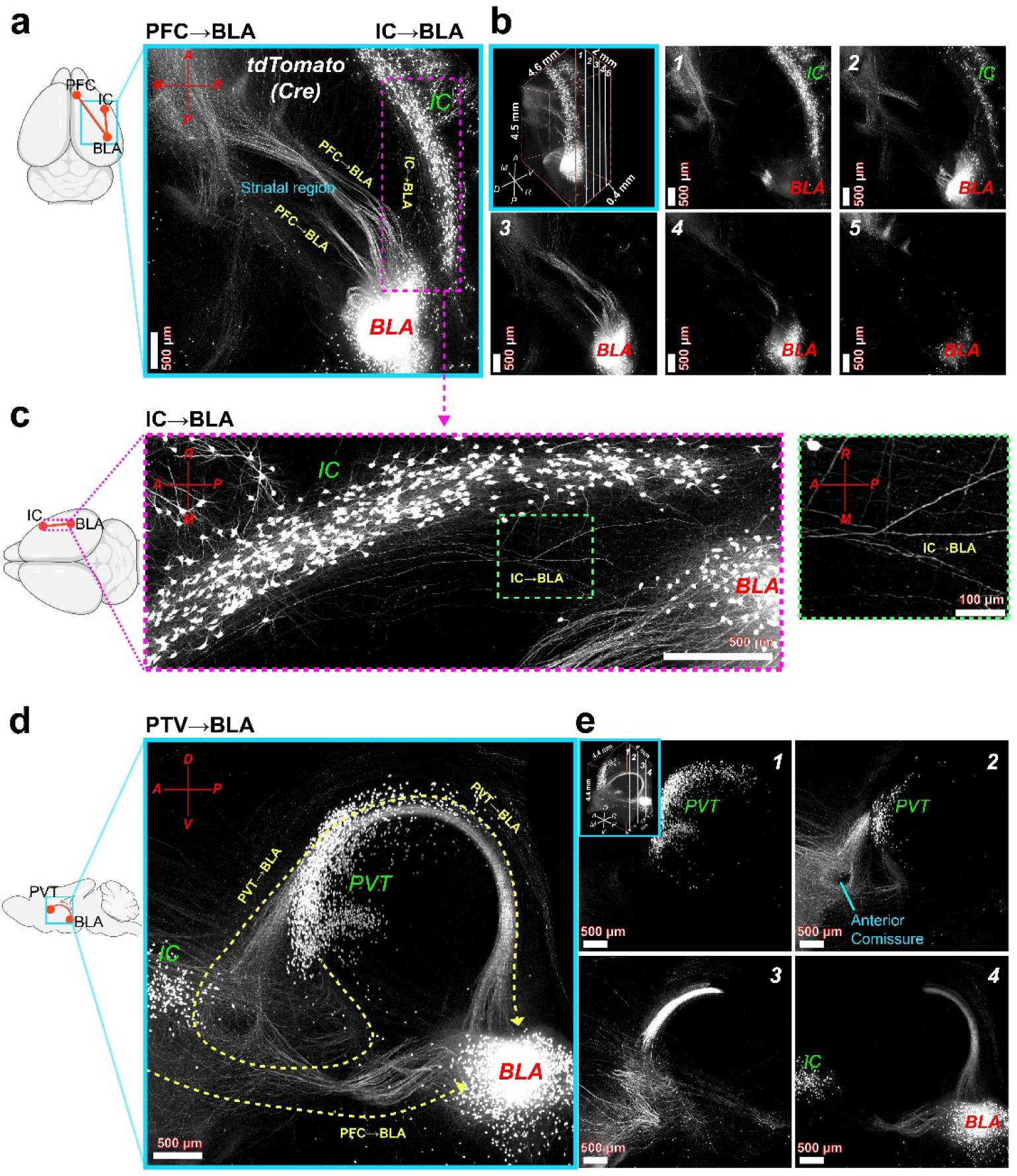
Detailed observation of axonal connections within the MNM centered on the PFC→BLA circuit (MNM^PFC→BLA^) (a) Maximum intensity projection (MIP: 2 mm) from a stack of virtual horizontal images showing axonal projections from PFC to BLA and IC to BLA. tdTomato signals are converted to white to emphasize signals from axonal connections. (b) Segmented view of the horizontal stack in (a), presented from dorsal to ventral (MIP: 0.4 mm/ each segment). (c) Left, magnified image of IC to BLA connections (MIP: 2 mm); right, further magnification. (d) Maximum intensity projection (4 mm) from a stack of sagittal section illustrating axonal projections from PVT to BLA and PFC to BLA. (e) Sequential sagittal sections from (d) shown from medial to lateral (MIPs: 0.6, 1, 1.6, 1.2 mm). Note: Directional markers indicate the following orientation: A (anterior), P (posterior), L (left), R (right), M (medial), D (dorsal), and V (ventral).

Recognizing the limitations in selectivity of conventional rAAV tracers, we developed the MNM-specific tracing method (Fig. 1e). This technique is an improvement of our previously established strategy for identifying single long-range neural circuits^36–39^ (see Extended Fig. 3 and Video 3). We utilized rAAV2/1 and rAAV2.retro serotypes that are engineered to bidirectionally deliver N-terminal and C-terminal Cre components from soma and axonal terminal regions, respectively^41^. Specifically, we created anterograde rAAV2/1-Ef1α-*CreN-InteinN* and retrograde rAAV2.retro-Ef1α-*InteinC-CreC*. Initially, we validated the successful Intein-mediated Cre reconstitution via co-expression of both rAAVs *in vitro* (Extended Fig. 4). Figure 1e depicts the *in vivo* mechanism of MNM approach. Upon injection into two distinct brain regions, these vectors are designed to travel along the axons of circuit neurons connecting those regions, undergoing reconstitution into full-length Cre. Furthermore, the trans-synaptic transport ability of rAAV2/1 ensures that Cre reconstitution also occurs in postsynaptic neurons of intermediary hub regions, which indirectly connect the two injection sites, thereby forming an MNM.

**Figure 3.**
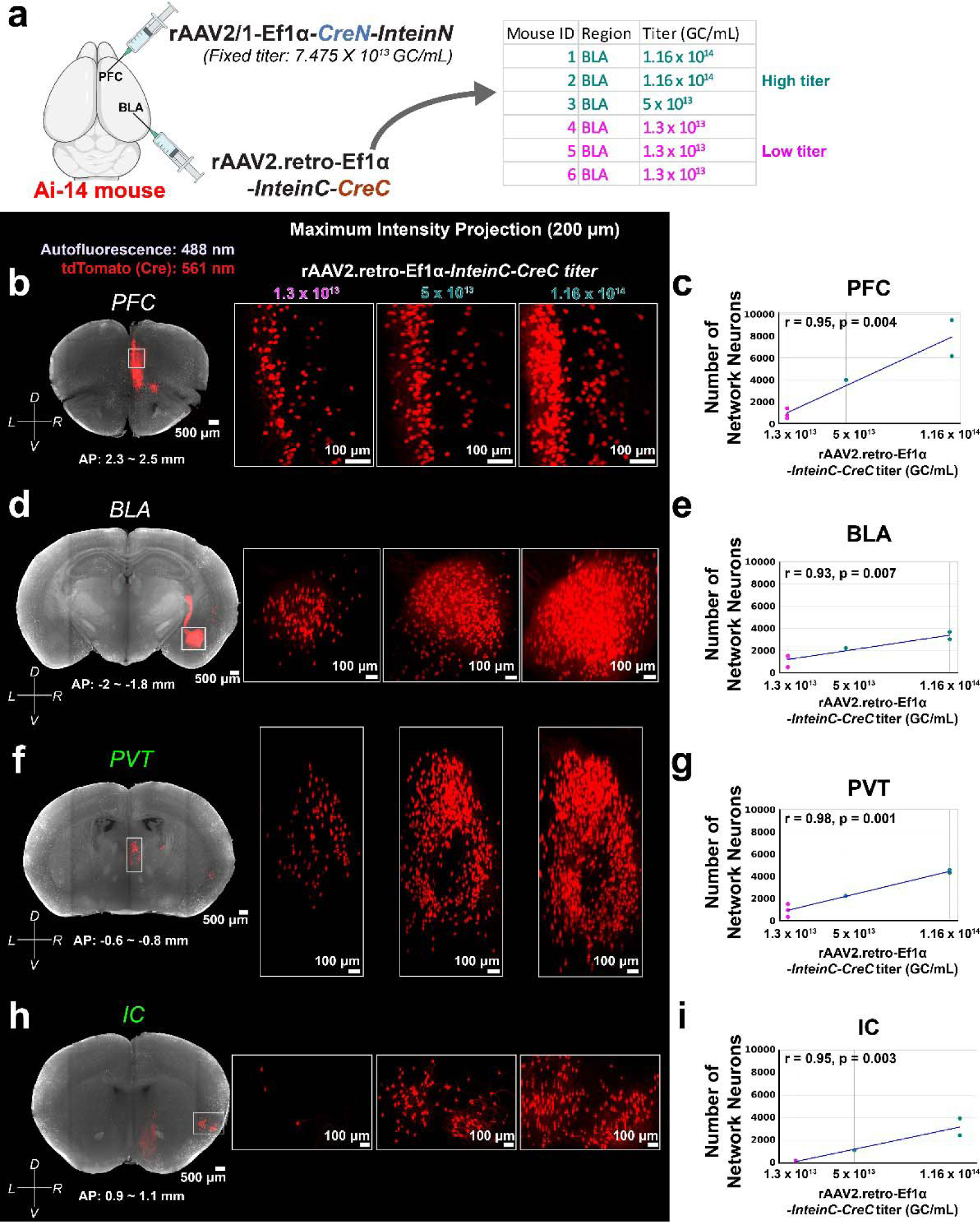
Relationship between rAAV titers and efficiency of MNM tracing. (a) Schematic of virus injection and the table of the varied rAAV titer conditions in MNM tracing for all six mice. For all animals, the titer of rAAV2/1-Ef1α*-CreN-InteinN* injected in the PFC was consistently 7.475×10^13^ GC/mL. However, the titer of rAAV2.retro-Ef1α*-InteinC-CreC* injected in the BLA varied, ranging from the lowest at 1.3×10^13^ GC/mL to 5×10^13^ or 1.16×10^14^ GC/mL. (b) Visualization of network neurons in the PFC: (Left) A virtual coronal section of the PFC from a mouse with rAAV2.retro-Ef1α*-InteinC-CreC* at a titer of 5×10^13^ GC/mL. (Right) Magnified images of PFC network neurons across varying titer levels for rAAV2.retro-Ef1α*-InteinC-CreC*, ranging from 1.3×10^13^ to 5×10^13^ and 1.16×10^14^ GC/mL. Each image is a maximum intensity projection (MIP) from a 200 µm stack. (c) Correlation analysis of the PFC network neuron expression with rAAV2.retro-Ef1α*-InteinC-CreC* titer. (d) Visualization of network neurons in the BLA (MIP: 200 µm). (e) Correlation analysis of the BLA network neuron expression with rAAV2.retro-Ef1α*-InteinC-CreC* titer. (f) Visualization of network neurons in the PVT (MIP: 200 µm). (g) Correlation analysis of the PVT network neuron expression with rAAV2.retro-Ef1α*-InteinC-CreC* titer. (h) Visualization of network neurons in the IC (MIP: 200 µm). (i) Correlation analysis of the IC network neuron expression with rAAV2.retro-Ef1α*-InteinC-CreC* titer. Note: lines in c, e, g, and i indicates least squares error. Note: Directional markers indicate the following orientation: L (left), R (right), D (dorsal), and V (ventral).

**Figure 4.**
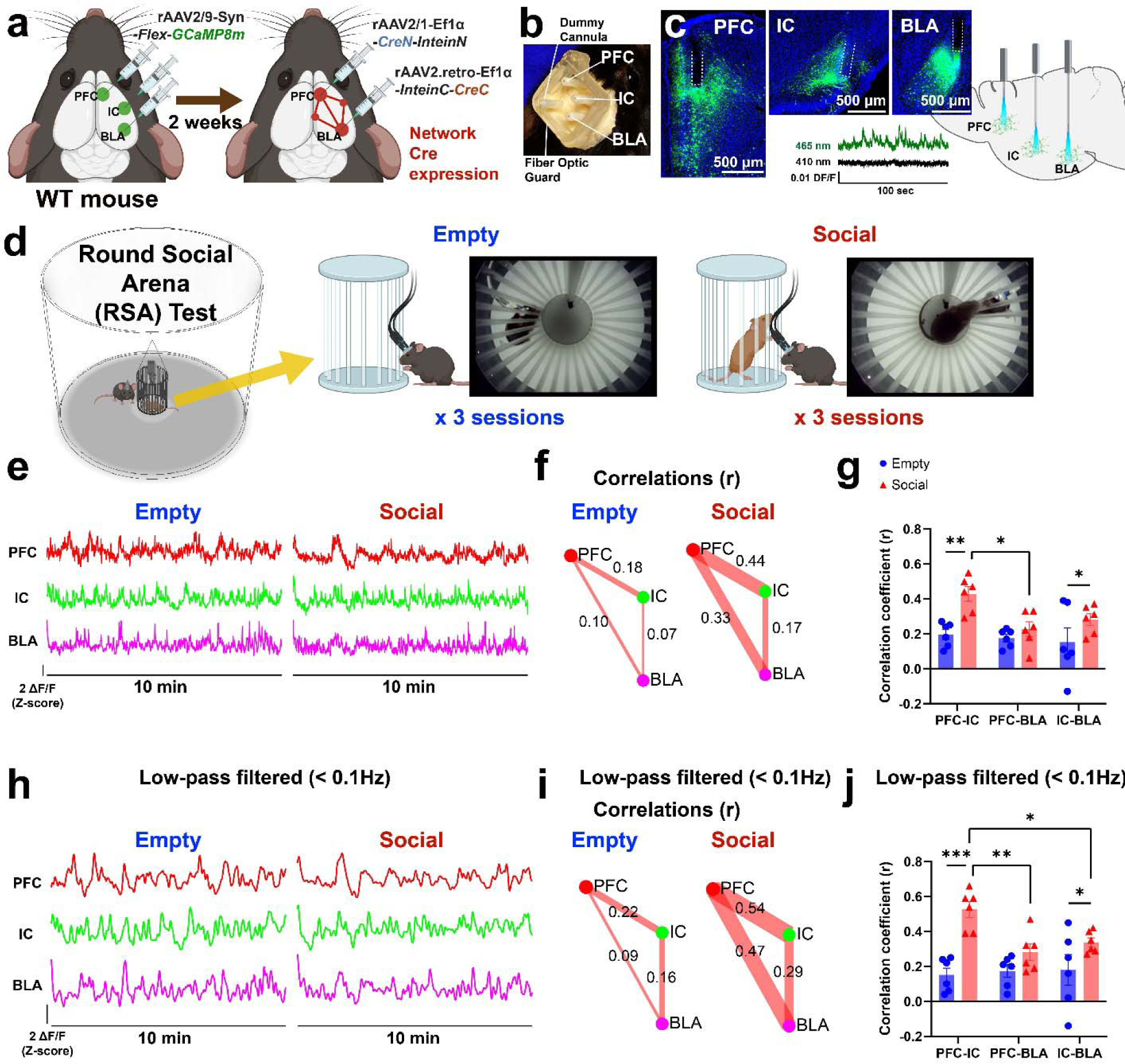
Monitoring network activity of MNM^PFC→BLA^ during social context. (a) A schematic illustrating the serial virus injection protocol for MNM-specific expression of GCaMP, a genetically encoded calcium indicator. Initially, rAAV2/9-Syn*-Flex-jGCaMP8m* is injected into the right hemisphere’s PFC, IC, and BLA of wild-type (WT) mice. Following a two-week interval, rAAV2/1-Ef1α*-CreN-InteinN* is injected into the PFC and rAAV2.retro-Ef1α*-InteinC-CreC* into the BLA, both in the right hemisphere. (b) An example image of the fiber optic cannula stubs implanted in the PFC, IC, and BLA of right hemisphere for fiber photometry recording. An extra dummy stub was cemented above the left hemisphere to secure the connection of extra branch from a 4-way fiber optic patch cable. A 3d-printed fiber guard was cemented around the stubs for protection. (c) (Left) Histological images displaying GCaMP8m expression and fiber-optic implant traces in the brain, with 500 µm scale bars. (Bottom) Example fiber photometry signals from the IC region, with neural activity indicated by fluorescence excited at 465 nm, representative of Ca^2+^ binding, and a control signal at 410 nm for isobestic correction. (Right) A schematic illustration detailing the simultaneous fiber photometry recordings from the network neurons in the PFC, IC, and BLA. (d) A schematic illustration of round social arena (RSA) setup used for studying social behaviors. Sessions were categorized as either ‘Empty’ or ‘Social’ based on whether a social stimulus mouse (female C3H/HeJ, aged P30-P50) was placed in the RSA’s inner cage. Each session, lasting 10 minutes, was part of a series of six sessions conducted over three days, comprising three Empty and three Social sessions. (e) Representative fiber photometry signals from PFC, IC, and BLA, corrected using isobestic signals, recorded during ‘Empty’ and ‘Social’ sessions. (f) Correlations between the fiber photometry signals of the three node regions (PFC, IC, BLA) during representative Empty and Social sessions, displayed in a network graph format. The thickness of each edge in the graph corresponds to the strength of the correlation coefficient. (g) Bar graphs comparing network correlation coefficients across Empty and Social sessions. The correlations in PFC-IC pair and IC-BLA pair are significantly heightened during Social sessions compared to Empty ones (p < 0.05). Notably, PFC-IC correlations during Social sessions are significantly stronger than PFC-BLA correlations in the same context (p < 0.05). (h) Sub-0.1 Hz photometry signals at representative Empty and Social sessions obtained by applying low-pass filter (<0.1 Hz) to fiber photometry signals in (g). (i) Sub-0.1 Hz network correlations at representative Empty and Social sessions. (j) Bar graphs comparing sub-0.1 Hz network correlations coefficients across Empty and Social sessions. The correlations in PFC-IC pair and IC-BLA pair are significantly heightened during Social sessions compared to Empty ones (p < 0.01). Notably, PFC-IC correlations during Social sessions are significantly stronger than PFC-BLA and IC-BLA correlations in the same context (p < 0.05). All bar graphs represent mean ± SEM. Asterisks indicate statistically significant p-values (*: p < 0.05, **: p < 0.01, ***: p < 0.001).

To demonstrate the effectiveness of MNM tracing, we injected rAAV2/1-Ef1α-*CreN-InteinN* into the right PFC and rAAV2.retro-Ef1α-*InteinC-CreC* into the right BLA of Ai-14 mice. Imaging of the cleared brains revealed dense tdTomato signals predominantly in key node regions of the ipsilateral hemisphere to the injections sites, including the PFC and BLA, and hub regions such as the paraventricular nucleus of thalamus (PVT) and insular cortex (IC) (Fig. 1f-g, Extended Fig. 5a, Video 4). These regions collectively formed the MNM^PFC→BLA^. We observed detailed neuronal cell bodies and axonal connections within this network (Fig. 1g). Additionally, extra hub nodes with smaller neuron populations were identified in the posterior part of the ipsilateral hemisphere, including areas such as the temporal association areas (TEa), ectorhinal areas (ECT), perirhinal areas (PERI), entorhinal areas (ENT), posterior piriform region (pPIR), and postpiriform transition areas (TR) (Fig. 1h). Statistical analysis confirmed significantly higher expressions of network neurons in the ipsilateral PFC, BLA, PVT, IC, and the combined TEa/ECT/PERI regions compared to the contralateral hemisphere (Two-way ANOVA with repeated measures, Šídák post-hoc test, p < 0.05) (Extended Fig. 5b). In addition, we observed extra axonal projections to the cerebral peduncle (cpd) (Extended Fig. 6). Notably, tdTomato signals found in cpd region were mostly from axons and not from neuronal cell bodies, suggesting that these may be collateral projections of the _MNM_PFC_→_BLA.

### Anatomically verifiable connectivity within the MNM^PFC→BLA^

MNM tracing not only highlighted the neurons at the network nodes but also delineated the axonal connections between them, especially the nodes including PFC, BLA, PVT, and IC. Detailed examination of the 3-D cleared brain images revealed a distinct bundle of axons projecting from the PFC directly to the BLA (Fig. 2a). The pathway originated in the PFC, coursed diagonally through the striatal regions (Extended Fig. 7a), and terminated at the BLA. Virtually reconstructed serial horizontal brain sections from superior to inferior planes further clarified the PFC→BLA trajectory, distinguishing it from adjacent pathways (Fig. 2b and Video 4). In contrast, the pathways from the IC to the BLA were more dispersed throughout the temporal lobe and did not form a dense bundle, yet they were still traceable (Fig. 2c).

The pathway from the PVT to the BLA displayed a complex route, with its intricate path discernible only through 3-dimensional analysis (Fig. 2d and Video 4). A virtually reconstructed sagittal section revealed that the PVT→BLA axons stray from a linear trajectory; initially projecting anteriorly, they navigate around the anterior commissure, then follow a curved bundle of axons that rises and falls across the hemisphere before reaching the BLA (Fig. 2d, Extended Fig. 7b). Serial sagittal sections further delineated this elaborate route (Fig. 2e).

### Optimization of MNM tracing with varying levels of virus titers

In our initial MNM tracing experiment, we utilized a fixed titer for rAAV2/1-Ef1α-*CreN-InteinN* (7.475×10^13^ GC/mL) and two different titers for rAAV2.retro-Ef1α-*InteinC-CreC* (5×10^13^ and 1.16×10^14^ GC/mL) (Extended Fig. 5a). Given that split-Cre reconstitution relies on the probability of two rAAV vectors infecting the same neuron, we hypothesized that titer levels might significantly influence MNM tracing efficiency. To test this, we conducted another MNM tracing experiment with a reduced titer of rAAV2.retro-Ef1α-*InteinC-CreC* (1.3×10^13^ GC/mL) across three mice (Fig. 3a, Extended Fig. 8a). This adjustment resulted in an overall reduction in MNM signals (Extended Fig. 8b). A detailed analysis revealed strong positive correlations (Pearson’s, r > 0.9) between the rAAV titers and the number of Cre-expressing neurons in key network nodes, including the PFC, BLA, PVT, and IC (Fig. 3b-i). These correlations were all statistically significant (p < 0.05). Notably, under the lowest titer condition, the IC showed a drastic reduction in network neurons almost to zero (Fig. 3h-i). Similar positive correlations were also observed in all posterior hub regions (Extended Fig. 9). These findings suggest that an optimized titer configuration is crucial for effective MNM tracing.

For MNM^PFC→BLA^, our findings indicate that optimal titers are above 7.475×10^13^ GC/mL for rAAV2/1-Ef1α-*CreN-InteinN* and above 5×10^13^ for rAAV2.retro-Ef1α-*InteinC-CreC*, respectively. To extend these findings, future research exploring a variety of MNMs will be essential to determine the most effective titer configurations for each specific network module.

### Monitoring MNM activity using Cre-dependent Ca^2+^ indicator and fiber photometry

To monitor the activity of MNM^PFC→BLA^, we utilized a Cre-dependent Ca^2+^ indicator, rAAV2/9-Syn*-Flex-jGCaMP8m*, coupled with fiber photometry imaging to monitor populational neuronal activity. Three network node regions were chosen for fiber photometry recording based on the available spatial interval that allowed for simultaneous recording: the PFC, IC, and BLA of the right hemisphere. Initially, we injected rAAV2/9-Syn*-Flex-jGCaMP8m* (5.78×10^13^ GC/mL) into the right hemisphere’s PFC, IC, and BLA of WT mice (n=2) (Fig. 4a). After a two-week incubation period, we administered high-titer rAAV2/1-Ef1α-*CreN-InteinN* (6.35×10^14^ GC/mL) to the PFC and rAAV2.retro-Ef1α-*InteinC-CreC* (3.29×10^14^ GC/mL) to the BLA. One month following these injections, we implanted fiberoptic cannula stubs into the three network node regions: PFC, IC, and BLA, to monitor the populational neural activity from these node regions (Fig. 4b-c).

In conjunction with fiber photometry, we utilized the Round Social Arena (RSA), a tool we previously developed for monitoring social interaction^44^ (Fig. 4d; Video 5). The RSA features a circular arena with a cage at the center. During ‘Empty’ sessions, this cage remained empty, serving as a control. In contrast, during ‘Social’ sessions, it housed a social stimulus mouse (C3H/HeJ, female, aged P30-P50). In total, each mouse was subjected to 6 sessions (3 Empty, 3 Social).

Figure 4e presents fiber photometry recordings from the three node regions – PFC, IC, and BLA – during example Empty and Social sessions. Utilizing these recordings, we analyzed the correlation coefficients between each pair of nodes (Fig. 4f). A two-way ANOVA with repeated measures indicated a significant impact of the social stimulus on these correlations [F(1, 15) = 17.34, p < 0.001], with an observed increase in inter-regional correlations during Social sessions (Fig. 4g). Subsequent Šídák post-hoc tests indicated a marked rise in PFC-IC and IC-BLA correlations during Social sessions compared to Empty ones (p < 0.05), but no significant change in PFC-BLA correlations (p = 0.38). Notably, the PFC-IC correlations were significantly stronger than PFC-BLA correlations during Social sessions (p < 0.05). Further analysis using fast-fourier transform (FFT) on the fiber photometry signals revealed peak signal magnitudes in the sub-0.1 Hz range (Extended Fig. 10a). Consequently, we applied a low-pass filter to the photometry signals to isolate the sub-0.1 Hz components (Fig. 4h) and re-evaluated the correlations (Fig. 4i). This analysis demonstrated a significant increase in correlations in the sub-0.1 Hz range during Social sessions compared to Empty ones, specifically for the PFC-IC and IC-BLA pairs [two-way ANOVA with repeated measures; F(1, 15) = 25.42, p < 0.001; Šídák post-hoc tests, p < 0.05] (Fig. 4j). Moreover, the PFC-IC correlation during Social sessions at sub-0.1 Hz was significantly higher than PFC-BLA and IC-BLA correlations (p < 0.05). Analysis of correlations in other frequency ranges, specifically those greater than 1 Hz and between 0.1 to 1 Hz, revealed no significant changes in the strength of node pair correlations during Social sessions compared to Empty ones (Extended Fig. 10b-g). These data suggest that alterations in correlative neural activity during social contexts predominantly occur within the sub-0.1 Hz frequency component.

This may reflect the slow oscillatory synchrony that is utilized for communication within distal networks^45^. Similarly, multi-regional synchrony at sub-0.1 Hz has been consistently reported in human resting state fMRI, which was suggested as baseline functional connectivity^46–48^.

In summary, we successfully monitored the three key nodes of the MNM^PFC→BLA^, demonstrating specific changes in network activity in response to social contexts. This approach provides a proof of concept for observing detailed modular network functions during behavior with anatomical precision.

## Discussion

In this study, we introduce the MNM approach, a significant advancement in analyzing brain subnetworks. This method allows for the precise targeting of specific modular networks composed of monosynaptic connections, a crucial capability that was previously unattainable with conventional rAAV tracers.

The MNM approach can serve as a versatile platform that integrates various advanced techniques in neuroscience, such as Cre-dependent conditional chemogenetics^49, 50^ and optogenetics^51–53^. These combined approaches enable selective manipulations of neurons in the network modules, empowering researchers to investigate the neural network mechanisms underlying specific behaviors.

Another pivotal aspect of MNM approach is exploring the role of neuropsychiatric risk-genes within modular networks. Our previous studies, which conducted knockout of *Shank3* or *ArpC3* genes in the PFC→BLA circuit, demonstrated their impacts on social behavior^36, 37^. Extending these efforts to the MNM level will deepen our understanding of the pathogenic network mechanisms underlying neuropsychiatric disorders. In the future studies, we can examine the functions of various risk-genes within the MNM framework using conditional transgenic mouse models.

Moreover, by extending MNM mapping to multiple interconnected modules, we can build a more comprehensive brain connectome atlas, composed of smaller modular networks. The module-focused atlas will provide novel insights into the whole-brain-level functional connectome. In this framework, researchers can construct a comprehensive database of modular network activity profiles associated with specific behaviors using MNM-specific multi-fiber photometry, as demonstrated in our study. Furthermore, this database can be expanded through integration of the functional datasets from MNM-specific chemogenetics and optogenetics, along with data from MNM-specific transgenic mouse models associated with neuropsychiatric disorders. In addition, network-specific electrophysiological examinations are achievable by targeting the fluorescence-labeled node neurons in the network. Finally, the MNM approach has the potential to significantly complement fMRI-based human brain network studies. Utilizing the MNM database as a reference, fMRI studies can focus on specific target brain regions to investigate various network activities and related behaviors. Conversely, well-established human brain networks identified in fMRI studies, such as the default mode or salience networks^48, 54, 55^, could guide the application of MNM approach in animals. This collaboration enables in-depth analyses of brain networks, aiding in the understanding of their structure and functional roles in various behaviors. Taken together, this multifaceted repository can combine structural, functional, and genetic datasets from both normal and disease conditions, establishing a robust platform for data-mining studies utilizing machine learning methods.

An important consideration in our study was to address the limitations of current rAAV technology. While rAAV2/1 offers robust anterograde trans-synaptic transfer capabilities, it faces constraints, including potential for minor retrograde transport and limited anterograde transfer to specific neuron types such as serotonergic, cholinergic, and noradrenergic neurons^34^. In this regard, there may be potential limitations in our MNM tracing method. For instance, the current MNM tracing may include a little population of hub neurons that project to both the PFC and BLA, rather than only relaying inputs from the PFC to the BLA in one direction. Additionally, the MNM tracing may not adequately identify certain hub neurons that are serotonergic, cholinergic, or noradrenergic. Despite these limitations, rAAV2/1 remains the preferred choice for MNM’s component due to its pronounced anterograde bias and robust trans-synaptic transfer capabilities, as demonstrated in extensive anterograde tracing projects like the Allen Mouse Brain Connectivity Atlas^35^. Comparatively, other serotypes known for anterograde tracing, such as rAAV2/9 and rAAV2/8, also exhibit retrograde transport^56–59^ and have shown less effective anterograde trans-synaptic transfer than rAAV2/1^32^. Thus, future developments in rAAV technology, including the engineering of more specialized serotypes with fewer off-target effects, will be crucial for enhancing the precision and efficacy of the MNM approach.

In conclusion, the MNM approach offers a new scope in network neuroscience that bridges a critical gap between structural and functional connectivity methods. This technique has the potential to enhance our understanding of neural network topography and functions, particularly in the context of understanding detailed, even subtle changes in neural networks that underlie complex behaviors in both normal and disease conditions. Continued exploration and refinement of this method will be crucial in further unraveling the complexities of brain function and in developing targeted therapeutic strategies for a range of neuropsychiatric disorders.

## Methods

### Animals

This study utilized Ai-14 (B6;129S6-Gt(ROSA)^26Sortm14(CAG-tdTomato)Hze^/J; stock no. 007908), C57BL/6J (wild-type; stock no. 000664), and C3H/HeJ (stock no. 000659) mice, all procured from Jackson Laboratory. The genetic background of all the transgenic mice was 129Sv × C57BL/6J.

Ai-14 mice, aged 8 to 10 weeks at the time of virus injection, included both males and females. These mice, which express tdTomato upon Cre recombination, were utilized for histological analysis. They were housed in groups of 3 to 4 mice per cage.

C57BL/6J wild-type (WT) mice, also aged 8 to 10 weeks at the time of virus injection and 16 to 18 weeks during behavioral experiments, were all males. These mice were used for concurrent fiber photometry recordings and behavioral experiments. They were pair-housed with another WT mouse. C3H/HeJ mice, utilized as social stimuli, were females aged between postnatal day 30 to 50.

All mice were housed in the Laboratory Animal Care Unit (LACU) of the University of Tennessee Health Science Center. Behavioral tests were conducted during the light cycle. All procedures adhered to protocols approved by the University of Tennessee Institutional Animal Care and Use Committee and complied with US National Institutes of Health guidelines.

### rAAV production

We have produced a series of recombinant adeno-associated viruses (rAAVs) for use in this study. These include rAAV2/1-hSyn-*Cre*, rAAV2.retro-hSyn-*Cre*, rAAV2/1-Ef1α-*CreN-InteinN*, rAAV2.retro-Ef1α-*InteinC-CreC*, rAAV2/8-Ef1α-*CreN-InteinN*, and rAAV2/9-Syn*-Flex-jGCaMP8m*. The production and subsequent titer measurement of these rAAVs were based on protocols from our previous studies^38, 60^, with necessary adjustments. Notably, to produce rAAV2/1-Ef1α-*CreN-InteinN* and rAAV2/1-hSyn-*Cre*, we utilized the pAAV2/1 packaging plasmid (Addgene #112862), diverging from our previous use of the pAAV 2/8 or 2/9 plasmids. Other plasmids for rAAV packaging were obtained from Addgene, including rAAV2-retro helper (Addgene #81070)^31^, pAAV2/8 (Addgene #112864), pAAV2/9 (Addgene #112865), pAAV-hSyn-*Cre* (Addgene #170367)^61^, and pGP-AAV-Syn*-Flex-jGCaMP8m-WPRE* (Addgene #162378)^62^, facilitating the production of our specific rAAVs.

### Validation of rAAV-mediated split-Cre reconstitution in vitro

HEK293T cells were seeded onto 12-well plate (2×10^5^ cells/ well). One day post-seeding, 2 µL of one or more viral vectors were administered to each well, according to specific treatment conditions. The viral vectors used, along with their respective titers, were as follows: rAAV2/8-Ef1α*-Flex-GFP* (2.3×10^13^ GC/mL), rAAV2/1-Ef1α-*CreN-InteinN* (6.35×10^14^ GC/mL), and rAAV2.retro-Ef1α-*InteinC-CreC* (7.43×10^14^ GC/mL).

After a two-day incubation period at 37°C, the cells were fixed with 4% paraformaldehyde for 10 min and treated with 4’,6-diamidino-2-phenylindole (DAPI; Sigma-Aldrich) diluted to a 1/10,000 concentration from the stock solution. This step was undertaken for nuclear staining to facilitate cell visualization. Finally, the wells were imaged using the CELENA-S Digital Imaging System Microscope (Logos Biosystem) with a 20x objective lens.

### Stereotaxic surgeries

Stereotaxic surgeries for viral injections were conducted in accordance with the protocols established in our previous studies^37, 38^. Briefly, mice aged 8-10 weeks were anesthetized using an isoflurane vaporizer (VetEquip). After shaving and cleaning their heads with betadine and 70% ethanol, the mice were securely positioned in a stereotaxic frame (David Kopf Instruments). A mixture of isoflurane vapor and oxygen was continuously delivered to the mice through a gas head holder (David Kopf Instruments) throughout the procedure. Craniotomies were performed at specific coordinates corresponding to the target brain regions. For virus delivery, we used a glass micropipette backfilled with mineral oil, attached to a Nanoject II injector (Drummond Scientific Company). The viral solutions were administered at a controlled rate over a 30-minute period directly into the target areas. To minimize the risk of backflow, the micropipette was left in place for at least 5 minutes after the injection. All viral injections were localized to the right cerebral hemisphere.

#### Anterograde tracing surgery from the PFC

To trace anterograde projections from the prefrontal cortex (PFC) and label recipient neurons, we administered 550 nL of rAAV2/1-hSyn*-Cre* (titer: 2.7×10^13^ GC/mL) into the PFC of an Ai-14 mouse (n=1). The injection coordinates were set to AP (anteroposterior): 2.5 mm, ML (mediolateral): 0.8 mm, and DV (dorsoventral): -1.5 to -1 mm from the brain surface.

#### Retrograde tracing surgery from the BLA

To trace regions projecting to the basolateral amygdala (BLA), we administered 150 nL of rAAV2.retro-hSyn*-Cre* (titer: 5.32×10^13^ GC/mL) into the BLA of an Ai-14 mouse (n=1). The injection coordinates were set as follows: AP: -1.8 mm, ML: 3.3 mm, and DV: -4.3 to -4 mm from the brain surface.

#### Single circuit tracing surgery for the PFC→BLA circuit

To selectively label neurons within the PFC→BLA circuit, we performed dual injections in Ai-14 mice (n=3). In the PFC, rAAV2/8-Ef1α*-CreN-InteinN* (500 nL; 1.1×10^13^ GC/mL) was injected at coordinates AP: 2.5 mm, ML: 0.8 mm, DV: -1.5 to -1 mm from the brain surface. Concurrently, the BLA was injected with rAAV2.retro-Ef1α*-InteinC-CreC* (200 nL; 1.51×10^13^ GC/mL) at AP: -1.8 mm, ML: 3.3 mm, DV: -4.3 to -4 mm.

#### MNM tracing surgery for the MNM^PFC→BLA^

To trace Monosynaptically-interconnected network module (MNM) centered on the PFC→BLA circuit with its additional hub regions, viral injections were performed in two batches of Ai-14 mice, with three mice in each batch. In the first batch, 600 nL of rAAV2/1-Ef1α-*CreN-InteinN* (7.475×10^13^ GC/mL) was injected into the PFC at AP: 2.5 mm, ML: 0.8 mm, DV: -1.5 to -1 mm from the brain surface. Concurrently, the BLA was injected with 200 nL of rAAV2.retro-Ef1α*-InteinC-CreC* at titers of either 5×10^13^ or 1.16×10^14^ GC/mL, at coordinates AP: -1.8 mm, ML: 3.3 mm, DV: -4.3 to -4 mm from the brain surface. For the second batch, the procedure was replicated, but the titer for rAAV2.retro-Ef1α-*InteinC-CreC* in the BLA was adjusted to 1.3×10^13^ GC/mL.

#### Surgery for selective GCaMP expression in the MNM^PFC→BLA^

To enable Ca^2+^ imaging of the MNM^PFC→BLA^, stereotaxic surgeries were conducted on two wild-type mice. Due to spatial constraints for multiple optic fiber implants in one hemisphere, we focused on three key nodes: the PFC, IC, and BLA. Initially, rAAV2/9-Syn*-Flex-jGCaMP8m* (5.78×10^13^ GC/mL) was administered into the right hemisphere at the following sites and volumes: PFC (AP: 2.5 mm, ML: 0.5 mm, DV: -1.5 to -1 mm; volume: 1000 nL), IC (AP: 0.8 mm, ML: 3.4 mm, DV: -2.8 to -2.3 mm; volume: 500 nL), and BLA (AP: -1.8 mm, ML: 3.3 mm, DV: -4.3 to -4 mm; volume: 400 nL). After two weeks, a second round of injections was performed, where rAAV2/1-Ef1α-*CreN-InteinN* (6.35×10^14^ GC/mL) into the PFC (AP: 2.5 mm, ML: 0.8 mm, DV: -1.5 to -1 mm; volume: 1000 nL) and rAAV2.retro-Ef1α-*InteinC-CreC* (3.29×10^14^ GC/mL) into the BLA (AP: -1.8 mm, ML: 3.3 mm, DV: -4.3 to -4 mm; volume: 400 nL).

#### Surgery for implanting multiple fiber optic cannulas

One month after viral injections, fiber optic cannula stubs (200 µm core diameter, 0.5 NA, 1.25 mm diameter ceramic ferrule; RWD Life Science R-FOC-L200C-50NA) were implanted into the wild-type mice. Holes were drilled at specific coordinates in the skulls for cannula placement: PFC (AP: 2.4 mm, ML: 0.3 mm), IC (AP: 0.8 mm, ML: 3.4 mm), and BLA (AP: -1.7 mm, ML: 3.5 mm). Additional holes were made in the left hemisphere for anchoring screws. Guide holes above the target regions were created using a blunt-cut 30G needle, with depths set at DV: -0.5 mm for PFC, -2.3 mm for IC, and -3.5 mm for BLA (measured from the brain surface).

During the cannula implantation, photometry signals were continuously monitored using a Multi-wavelength Fiber Photometry System (Plexon) to ensure optimal placement of the cannulas. Two lengths of cannulas were used: 3 mm for the PFC and 6.45 mm for the IC and BLA. The fibers were implanted to the following target depths: DVs of -1 to -0.8 mm for PFC, -2.3 mm for IC, and -3.8 to -3.7 mm for BLA. Each cannula was secured with C&B-Metabond (Parkell) cement, while the other holes were safeguarded using PBS-soaked cotton to prevent cement occlusion. For post-cementing of fiber optic cannula stubs, an additional dummy stub was cemented above the opposite hemisphere for secure connections with extra branch of the 4-way Plexon fiber optic cable. Lastly, a 3D-printed guard was placed around the cannulas for added protection.

### Perfusion and fixation of brain samples

Perfusion procedure follows our previous studies^63, 64^. Briefly, one month after the viral injections, which allowed for extensive tdTomato expression within the Monosynaptically-interconnected network, we initiated the perfusion process in the Ai-14 mice. The mice were deeply anesthetized using isoflurane and then underwent transcardial perfusion. This perfusion consisted of an initial flush with heparinized phosphate-buffered saline (PBS, 25 U/mL heparin) followed by fixation with 4% paraformaldehyde (PFA) in PBS. Subsequently, the brains were carefully extracted and placed in 4% PFA for overnight post-fixation at 4 °C. During this period, the samples were subjected to gentle shaking to ensure uniform fixation.

### Brain clearing

Following the perfusion and post-fixation, the brain samples were processed using the SHIELD (stabilization under harsh conditions via intramolecular epoxide linkages to prevent degradation) protocol developed by LifeCanvas Technologies^65^. This protocol entailed a two-step solution treatment, including SHIELD-off and SHIELD-on, designed to preserve endogenous fluorescence before initiating active tissue clearing. After the SHIELD treatment, the brains underwent active delipidation using the SmartClear Pro II device (LifeCanvas Technologies). This step is crucial for rendering the brain tissue optically transparent while maintaining the structural integrity and fluorescent signals of the tissue.

### Whole-brain imaging and analysis

Whole-brain imaging was conducted in the LifeCanvas Technologies. The brain samples underwent a preparation process for optical clarity and refractive index matching. Initially, the samples were incubated in 50% EasyIndex (Refractive Index, RI = 1.52, LifeCanvas Technologies) at 37 °C overnight. This step was followed by a 24-hour incubation in 100% EasyIndex to match the refractive index of the surrounding medium to that of the tissue. After the index matching process, imaging was performed using a SmartSPIM light sheet microscope (LifeCanvas Technologies). The microscope was equipped with a 3.6x objective lens [Numerical aperture (NA) = 0.2] and operated with a 4 µm step size for imaging. Dual-channel SPIM (Selective Plane Illumination Microscopy) imaging was conducted using 488 nm wavelength light for capturing autofluorescence and 561 nm for imaging tdTomato expression. This process yielded high-resolution image stacks with dimensions of 1.8 µm x 1.8 µm x 4 µm for each horizontal plane.

After acquiring whole-brain images, we processed and analyzed the images using SyGlass software (IstoVisio, RRID:SCR_017961), tailored for 3D reconstruction of biological images, for a detailed examination of the MNM^PFC→BLA^’s structural connectome in virtual reality. Using the Oculus Rift S headset and controllers, we analyzed complex axonal projections, such as the PVT→BLA pathway, which would have been challenging to discern with conventional confocal microscopy.

Subsequently, we used ImageJ Fiji (RRID:SCR_003070) for quantifying tdTomato-positive neurons. This process involved reslicing the z-stacks of horizontal plane images into coronal and sagittal planes. From these resliced planes, we generated maximum intensity projection (MIP) images with varying thicknesses (100 µm, 200 µm, and 400 µm). These MIP images allowed for the accurate counting of tdTomato-expressing neuronal cell bodies in each section. Furthermore, we utilized autofluorescence MIP images as a reference to delineate the regional boundaries of the network neurons, ensuring precise spatial localization of neuronal populations within the brain.

### Behavioral experiments

Two weeks following the fiber optic implantation in wild-type mice, we initiated behavioral experiments in parallel with fiber photometry recordings.

#### Round Social Arena (RSA) test

Expanding on our previous studies, the RSA test was specifically designed to emphasize social interactions while minimizing the distractors and spatial biases often encountered in traditional 3-chamber tests^37, 44^. The RSA consists of a circular arena (diameter: 49 cm) equipped with a central 3D-printed transparent bar cage (diameter: 8 cm, height: 10.5 cm) designed to house a social stimulus mouse. The cage features a cone-shaped roof to prevent climbing and includes a wide-angle (180°) fish-eye lens camera to facilitate detailed observation of the mice’s social behaviors.

The experimental protocol was structured over several days:

Habituation: Over three days, the animals were habituated to the empty arena for 15 minutes each day, allowing them to become accustomed to the new environment without the presence of a social stimulus.

RSA Test (Day 1): The test comprised three 10-minute stages, alternating between one Empty and two Social sessions, with 5-minute intervals between each session. During an Empty session, the mice explored the arena with an empty cage present, while in each Social session, a novel female C3H/HeJ mouse was introduced as a social stimulus.

RSA Test (Day 2): The test included two 10-minute stages – an Empty session followed by a Social session, separated by a 5-minute interval.

RSA Test (Day 3): The test concluded with another 10-minute Empty session.

### Fiber photometry recordings

Fiber photometry recordings of Ca^2+^ signals were performed using the Multi-Wavelength Fiber Photometry System (Plexon). A 2-meter patch fiber optic cable (OPT/PC-FC-4SLCF-100/110-2.0L/0.1L, Plexon), featuring four 10 cm branches at its end was utilized to simultaneously record photometry signals from the implanted fiber optics in the mouse brain. As recordings were taken from three channels (PFC, IC, BLA), the fourth branch of the cable was connected to a dummy cannula mounted on the mouse’s head.

Two distinct light wavelengths were employed for the recordings: 456 nm for monitoring Ca^2+^ signals, indicative of neural activity, and 410 nm to serve as an isobestic control. Before initiating the experiments, the light power at the tip of each fiber optic cannula (RWD Sciences) was meticulously measured using a power meter (PM100D, Thorlabs). The light intensities were calibrated to 30 µW for the 456 nm wavelength and 15 µW for the 410 nm wavelength. During the experiments, the Plexon Photometry system captured the emission responses elicited by these excitation lights, allowing for the recording of data for subsequent analysis.

### Analysis of the fiber photometry data

#### Preprocessing

For the preprocessing of the raw fiber photometry data, we employed pMAT^66^ and custom MATLAB (Mathworks, RRID:SCR_001622) scripts. The pMAT tool was used to correct the signals obtained from the 456 nm channel by subtracting the isobestic control data from the 410 nm channel. We then calculated the normalized change in fluorescence (ΔF/F) signals for the 456 nm channel. These corrected signals were subsequently temporally aligned with the behavioral data using a custom MATLAB script.

#### Computational analysis of fiber photometry data

The computational analysis of the fiber photometry signals was conducted entirely in MATLAB. We performed Fast Fourier Transform (FFT) on the normalized ΔF/F signals to determine the frequency ranges exhibiting maximum power. The FFT results revealed the highest signal magnitude in the sub-0.1 Hz range, with a decrease at higher frequencies, particularly in the 0.1-1 Hz range and further attenuation at frequencies over 1 Hz. Informed by these findings, we applied a series of filters to the data: a low-pass filter for frequencies below 0.1 Hz, a band-pass filter for the 0.1-1 Hz range, and a high-pass filter for frequencies above 1 Hz, all implemented using a Butterworth filter. Additionally, we conducted Pearson’s correlation analysis between each pair of nodes (PFC, IC, BLA) to assess the relationships in their activity. This correlation analysis provided insights into the synchronous or asynchronous nature of neural activity across these regions.

### Histology

Following the completion of fiber photometry imaging, the mice underwent perfusion, after which their brains were extracted and post-fixed in 4% paraformaldehyde (PFA) overnight at 4°C with gentle shaking. Subsequently, the brains were cryoprotected in 30% sucrose in phosphate-buffered saline (PBS) for three days to prepare for sectioning.

For histological examination, the brains were sectioned into 100 µm coronal slices using a Leica CM 1950 cryostat. These sections underwent a series of washes and staining procedures. Initially, they were washed in PBST (0.2% Triton X-100 in PBS) for 15 minutes to permeabilize the tissue. Following this, the sections were stained with DAPI (Sigma-Aldrich) to label the nuclei. After staining, the sections were washed three times in PBST to remove excess DAPI. This process enabled detailed visualization of neuronal structures and the distribution of GCaMP8m-positive neurons, as well as the traces of fiber optic implants.

The final step in the preparation process involved mounting the sections on slides and then coverslipping them with ProLong Glass anti-fade medium (Invitrogen) to preserve the fluorescence. Imaging of the sections was conducted using an LSM 710 confocal microscope (Zeiss) equipped with a 10× objective. The Zen software was utilized for capturing and analyzing the images.

### Statistical analysis

All statistical analyses were performed using Prism (Graphpad, RRID:SCR_002798) and MATLAB software. For the comparison of tdTomato expression and network neuron counts between hemispheres, we used two-way ANOVA with repeated measures, followed by Šídák post-hoc tests for specific pairwise comparisons. For fiber photometry data, Pearson’s correlation coefficient was calculated to assess relationships between neuronal activities in different network nodes. All tests were two-tailed, and a p-value of less than 0.05 was considered statistically significant. Results are presented as mean ± SEM. Comprehensive details of all statistical tests employed are provided in Supplementary Table 1.

## Reporting summary

Further information on research design is available in the Nature Research Reporting Summary linked to this article.

## Data availability

Further requests for data, resources, and reagents should be directed to and will be fulfilled by the lead contact, Il Hwan Kim.

## Code availability

Requests for custom MATLAB scripts used in this study should be directed to and will be fulfilled by the lead contact, Il Hwan Kim.

## Acknowledgement

This work was supported by NIH MH117429 and AG075000. We thank Drs. John Boughter, Matthew Ennis, and Lynn Dobrunz for their critical reading and comments, Dr. Fan Wang for providing split-Intein-Cre DNA sequences, and Leah (VanDenBosch) Chestnut at LifeCanvas Technologies for supporting light-sheet imaging. A subset of the illustrations was created with BioRender.com.

## Author contributions

SK and IHK designed this study. YK and IHK performed cloning. SK, YK, and YU packaged and purified rAAVs. SK and YU performed viral injection surgeries. YK conducted immunocytochemistry. SK performed perfusion, brain clearing, 3D-analysis of MNN structures, counting of network neurons, multi-fiber photometry imaging, behavioral tests, statistical analyses, and immunohistochemistry. This paper was written by SK and IHK and was edited by the other authors.

## Extended Figures

**Extended Figure 1.**
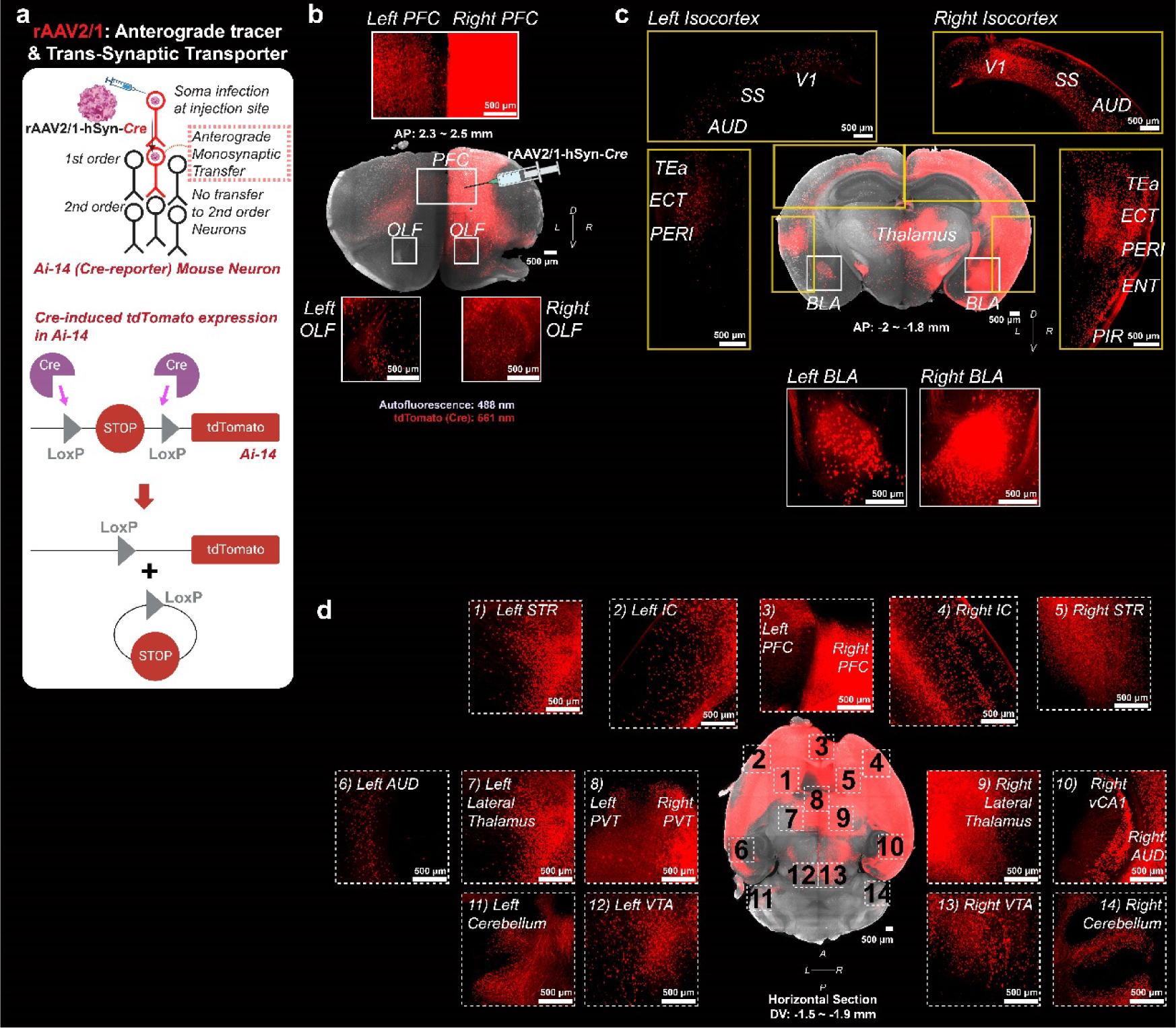
Anterograde tracing from the right PFC and its target neurons. (a) (Top) Schematic representation of the anterograde tracer rAAV2/1-hSyn-Cre, highlighting its journey to axonal projections and trans-synaptic transport to 1st-order postsynaptic neurons. (Bottom) Diagram illustrating the activation of tdTomato expression in Ai-14 transgenic mice by Cre recombinase, which removes a loxP-flanked stop codon, thereby initiating the transcription of the tdTomato sequence. (b) Virtual coronal section (AP: 2.3 ∼ 2.5 mm) from a cleared brain of an Ai-14 mouse injected with rAAV2/1-hSyn*-Cre* in the right prefrontal cortex (PFC), displaying tdTomato expression in the PFC and olfactory regions (OLF) across both hemispheres. (c) Virtual coronal section (AP: -2 ∼ -1.8 mm) displaying tdTomato expression in postsynaptic target neurons from the right PFC across various brain regions, including the basolateral amygdala (BLA), thalamus, temporal association areas (TEa), ectorhinal area (ECT), perirhinal area (PERI), visual area (V1), somatosensory areas (SS), and auditory areas (AUD) in both hemispheres. Notably, expression in the entorhinal area (ENT) and piriform area (PIR) is exclusive to the right hemisphere. (d) Virtual horizontal section (DV: -1.9 ∼ -1.5 mm from Bregma) revealing tdTomato expression in regions including the PFC, insular cortex (IC), striatum (STR), thalamus, paraventricular nucleus of the thalamus (PVT), AUD, ventral tegmental area (VTA), and cerebellum in both hemispheres, with exclusive expression in the ventral hippocampus CA1 (vCA1) of the right hemisphere. Note: Directional markers indicate the following orientation: A (anterior), P (posterior), L (left), R (right), D (dorsal), and V (ventral).

**Extended Figure 2.**
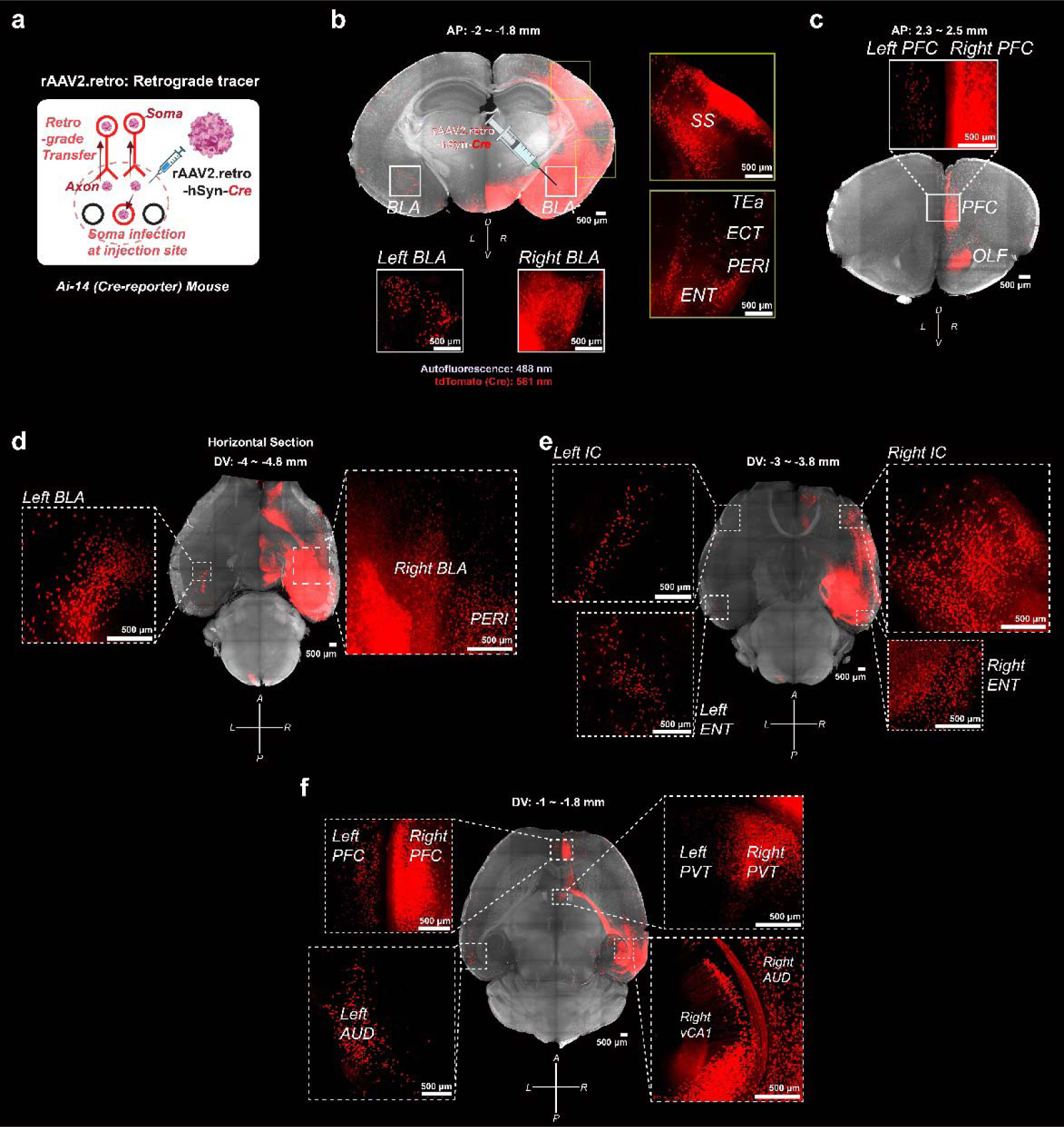
Retrograde tracing from the right BLA. (a) Schematic illustrating rAAV2.retro-hSyn-Cre’s retrograde travel property, in addition to its ability to infect neuronal cell bodies at the injection site. (b) Virtual coronal section (AP: -2 ∼ -1.8 mm) from an Ai-14 mouse, following injection of rAAV2.retro-hSyn*-Cre* into the right basolateral amygdala (BLA). This section shows tdTomato expression predominantly in the right BLA, ipsilateral to the injection site, with notable expression also observed in the left BLA. Enhanced tdTomato signals are evident in afferent regions of the right hemisphere, including the somatosensory area (SS), temporal association areas (TEa), ectorhinal area (ECT), perirhinal area (PERI), and entorhinal area (ENT). (c) Virtual coronal section (AP: 2.3 ∼ 2.5 mm) showing tdTomato expression in the anterior brain regions, with more pronounced expression in the right hemisphere’s prefrontal cortex (PFC) and exclusive signals in the right olfactory area (OLF). (d) Virtual horizontal section (DV: -4 ∼ -4.8 mm from Bregma) displaying tdTomato expression in the BLA of both hemispheres, stronger in the right. In this horizontal plane, unique tdTomato signals in temporal regions like the perirhinal area (PERI) are observed exclusively in the right hemisphere. (e) Virtual horizontal section (DV: -3 ∼ -3.8 mm from Bregma) illustrating tdTomato expression in the insular cortex (IC) and entorhinal area (ENT) of both hemispheres, with more intense signals in the right hemisphere. (f) Virtual horizontal section (DV: -1 ∼ -1.8 mm from Bregma) revealing tdTomato expression in the prefrontal cortex (PFC), paraventricular nucleus of the thalamus (PVT), and auditory areas (AUD) of both hemispheres, with stronger signals in the right. Notably, tdTomato signals in the ventral hippocampus CA1 (vCA1) are uniquely present in the right hemisphere in this section. Note: Directional markers indicate the following orientation: A (anterior), P (posterior), L (left), R (right), D (dorsal), and V (ventral).

**Extended Figure 3.**
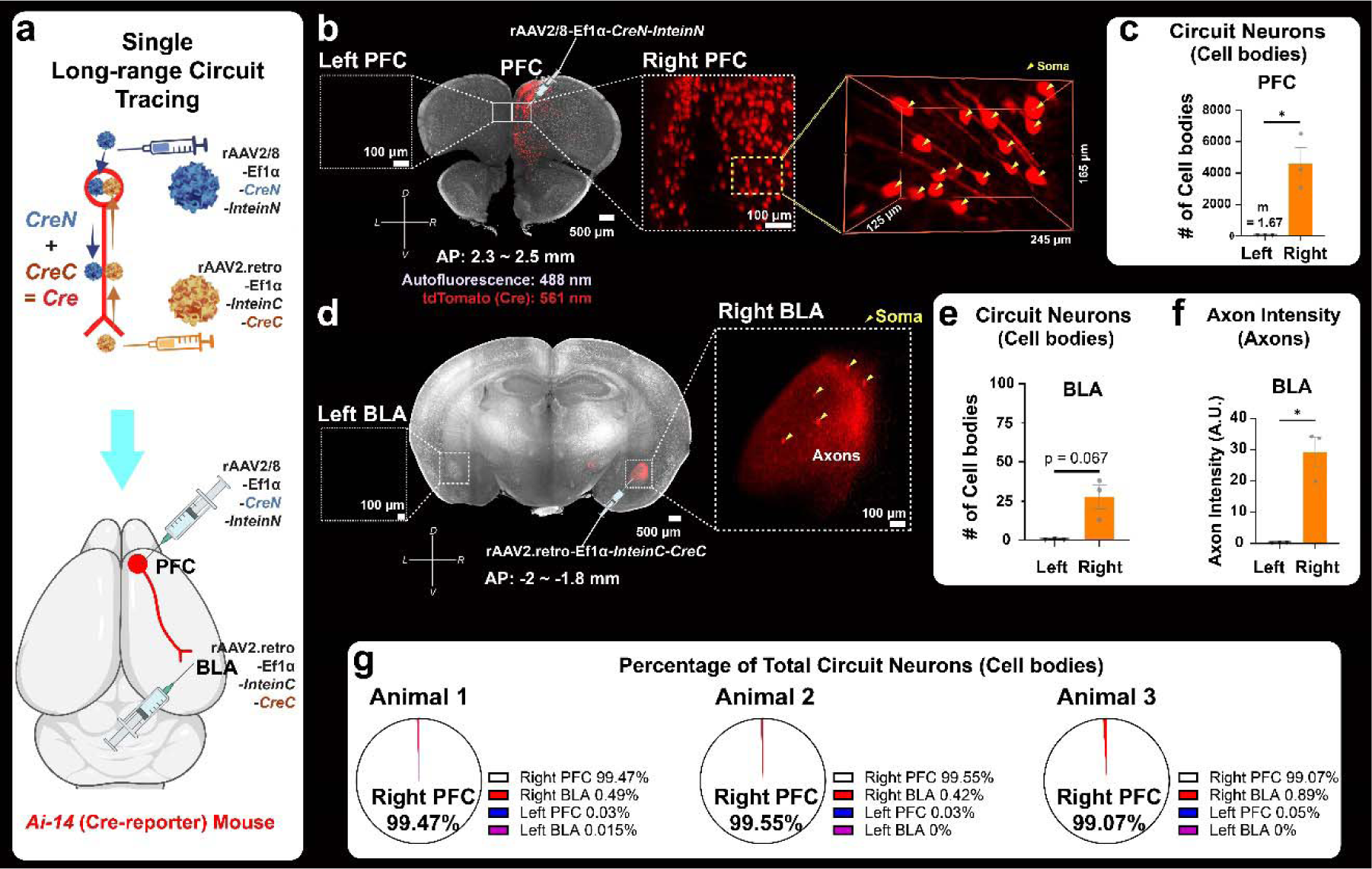
Tracing single long-range circuit: PFC→BLA circuit in the right hemisphere. (a) (Top) Schematic of single long-range circuit tracing. Anterograde rAAV2/8-Ef1α*-CreN-InteinN*, with minimal trans-synaptic property, is injected into neuronal cell bodies, while retrograde rAAV2.retro-Ef1α*-InteinC-CreC* targets axon terminals. CreN-InteinN and InteinC-CreC components converge within neurons, reconstituting a full-length Cre enzyme through split Intein-mediated reconstitution. (Bottom) Application of this technique to trace the PFC→BLA circuit in the right hemisphere. Anterograde rAAV2/8-Ef1α*-CreN-InteinN* is injected into the PFC and retrograde rAAV2.retro-Ef1α*-InteinC-CreC* is injected into the BLA of right hemisphere, enabling exclusive Cre reconstitution within the PFC→BLA circuit neurons. (b) (Left) Virtual coronal section (AP: 2.3 ∼ 2.5 mm) of cleared brain from an Ai-14 mouse with PFC→BLA circuit tracing in the right hemisphere. Circuit neurons labeled with tdTomato are almost exclusively found in the right PFC. (Right) Magnified image of the tdTomato-labeled circuit neurons in the right PFC. (c) Quantitative analysis of tdTomato-positive neuronal cell bodies in the PFC across three mice reveals an average of 4,616.67 labeled cells in the right PFC and only 1.67 in the left. A statistically significant difference was confirmed [paired t-test, t(2)=4.6, p < 0.05]. (d) (Left) Virtual coronal section (AP: -2 ∼ -1.8 mm) from a mid-posterior brain. Axon terminals expressing tdTomato signals are found exclusively in the right BLA. A few neuronal cell bodies in the right BLA also exhibit tdTomato expression, indicative of minimal trans-synaptic transport by rAAV2/8. (Right) A magnified image of the tdTomato signals in the right BLA. (e) Quantitative analysis of tdTomato-positive neuronal cell bodies in the BLA from three mice shows an average of 27.67 labeled cells in the right BLA and 0.33 in the left, with no significant statistical difference between hemispheres [paired t-test, t(2) = 3.671, p = 0.0669]. (f) Quantitative analysis of tdTomato-labeled axonal intensity in the BLA across three mice indicates an average intensity of 29.248 A.U. in the right BLA, significantly higher than 0.24 A.U. in the left BLA [paired t-test, t(2) = 6.084, p < 0.05], confirming effective isolation of the PFC→BLA circuit within the right hemisphere. (g) The percentage of total circuit neuron cell bodies found in each mouse demonstrates that over 99.07% of tdTomato-labeled circuit neuron cell bodies are located in the right PFC in all three mice. This finding confirms the successful isolation of the PFC→BLA circuit in the right hemisphere and shows the minimized trans-synaptic tagging of neurons receiving input from the circuit. All bar graphs represent mean ± SEM. An asterisk indicates a statistically significant p-value (*: p < 0.05). Note: Directional markers indicate the following orientation: L (left), R (right), D (dorsal), and V (ventral).

**Extended Figure 4.**
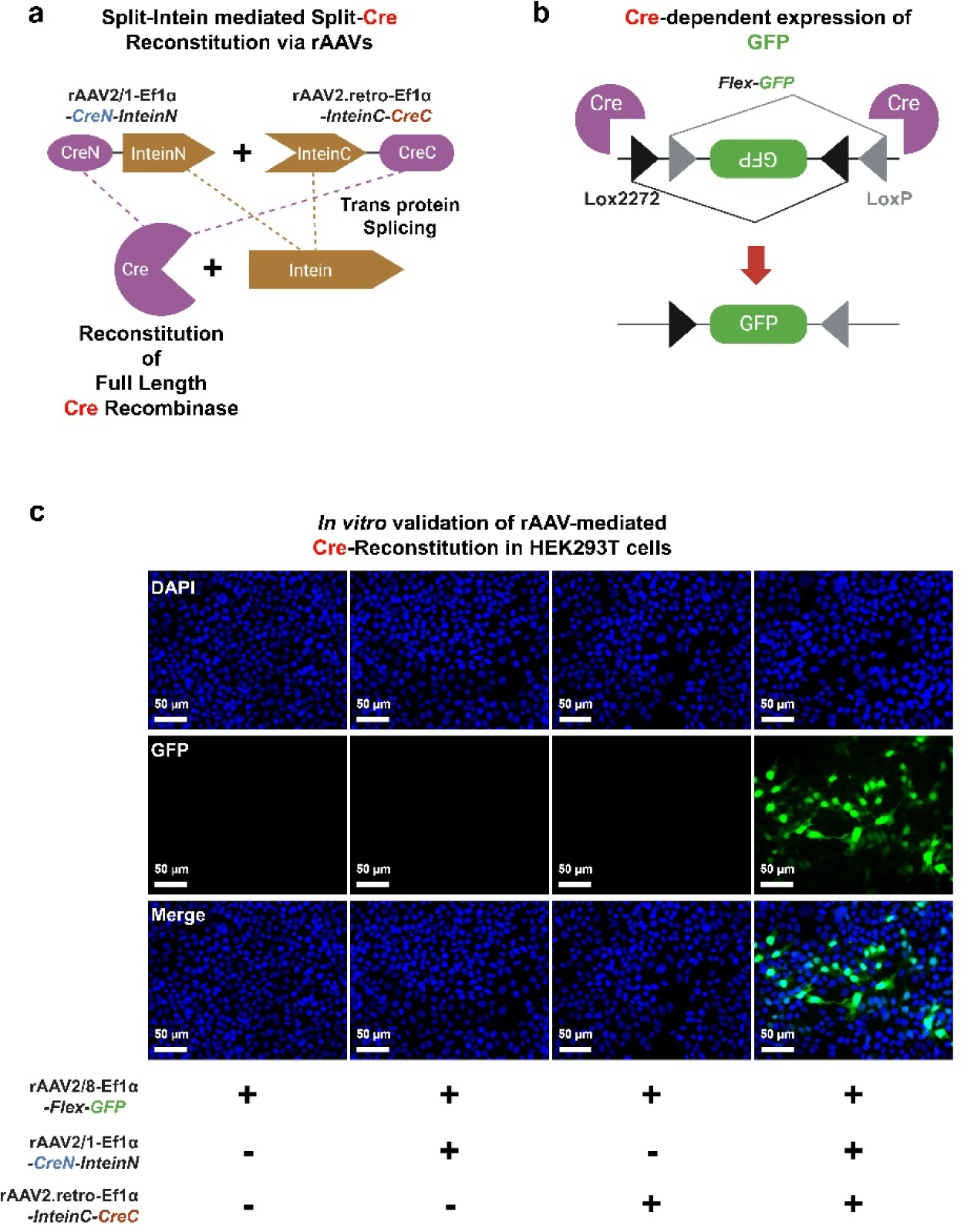
Mechanism of split-Intein mediated Cre reconstitution and its *in vitro* validation. (a) Diagram of split-Intein mediated split-Cre reconstitution. The vector rAAV2/1-Ef1α*-CreN-InteinN* carries the N-terminal portion of Cre recombinase (CreN) fused to the N-terminal portion of an Intein (InteinN), while rAAV2.retro-Ef1α*-InteinC-CreC* carries the C-terminal portion of an Intein (InteinC) fused to the C-terminal portion of Cre recombinase (CreC). When both vectors enter the same cell, the protein products from these vectors - the split-Cre segments fused with respective Intein segment - are synthesized. These protein parts then undergo a process where the Intein segments self-excise and ligate the CreN and CreC fragments, resulting in the reconstitution of a full-length, functional Cre recombinase. (b) Diagram of Cre-dependent expression of GFP (green fluorescent protein) using the Flex (FLEx: Flip-Excision)-GFP system. The Flex system employs a double-inverted orientation approach. Initially, a gene of interest—in this case, GFP is flanked by a pair of incompatible Lox sites: LoxP and Lox2272. These Lox sites are arranged in such a way that the GFP sequence is initially in a reverse orientation, making it non-functional. Upon the introduction of Cre recombinase, it recognizes these Lox sites and induces recombination between them. This recombination event flips the GFP sequence into the correct orientation, allowing for its expression. However, because LoxP and Lox2272 sites are incompatible for further recombination, the flip is irreversible, stabilizing the expression of GFP. This system ensures that GFP expression only occurs in cells where Cre recombinase is active, providing precise control over the gene expression. (c) *In vitro* testing of split-Cre reconstitution was successfully conducted in HEK293T cells using a combination of rAAV2/1-Ef1α*-CreN-InteinN*, rAAV2.retro-Ef1α*-InteinC-CreC*, and rAAV2/8-Ef1α*-Flex-*GFP vectors. GFP expression was observed only when all three vectors were co-introduced, confirming the successful reconstitution of Cre carried out by the rAAVs.

**Extended Figure 5.**
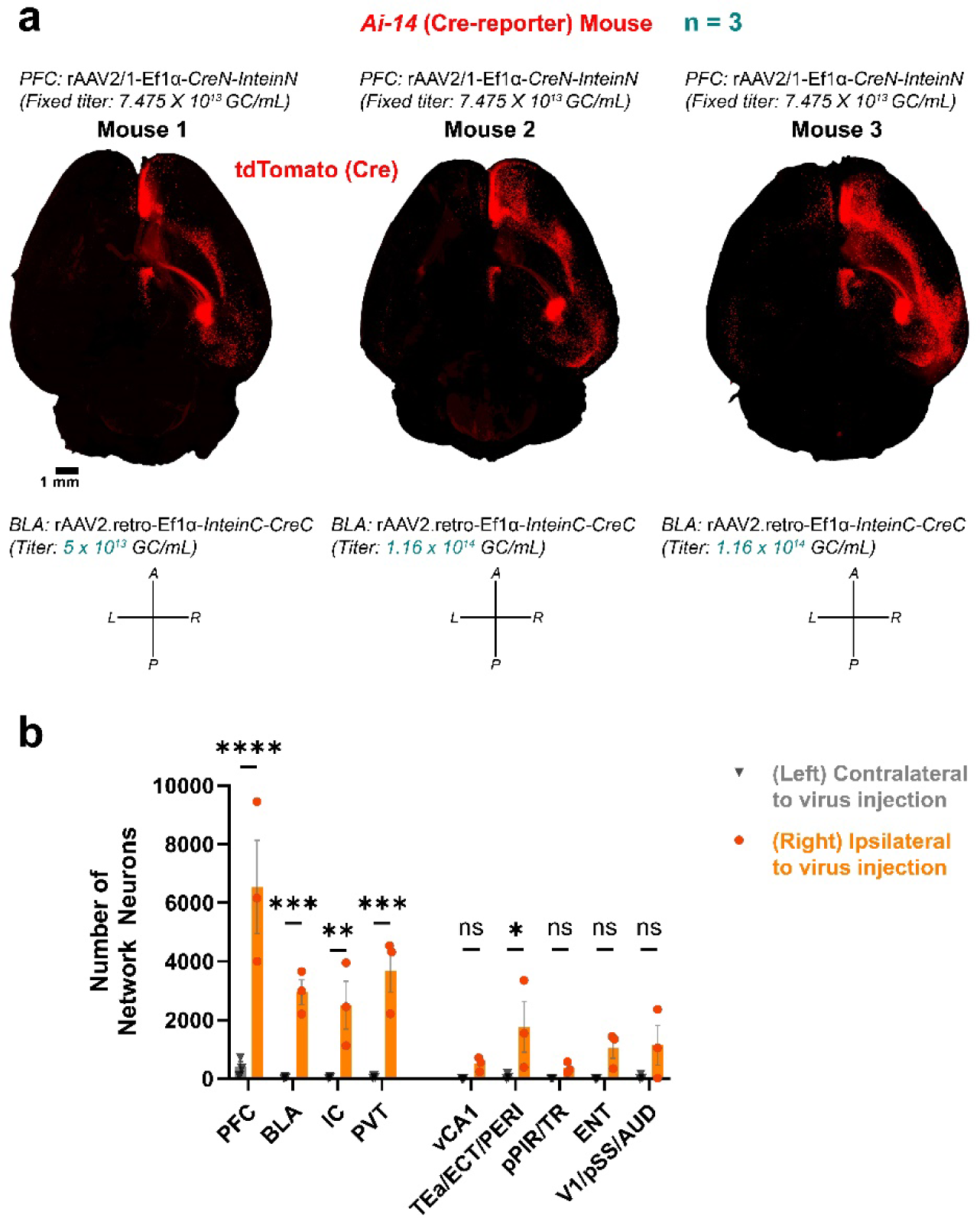
Cleared brain samples with MNM^PFC→BLA^ tracing. (a) Display of cleared brain samples from Ai-14 mice (n=3) with MNM^PFC→BLA^ tracing. In each sample, rAAV2/1-Ef1 *-CreN-InteinN* at a fixed titer of 7.475×10^13^ GC/mL was injected into the right PFC. The right BLA received varying titers of rAAV2.retro-Ef1α*-InteinC-CreC*: one mouse was injected with 5×10^13^ GC/mL and the other two with 1.16×10^14^ GC/mL. All samples all resulted in robust MNM^PFC→BLA^ signaling, with clear visualization of axonal connections across the brain. (b) Quantitative analysis of tdTomato-labeled network neurons across the two hemispheres in Ai-14 mice demonstrated a significant increase in expression in the right hemisphere (ipsilateral to virus injection) compared to the left (contralateral). A two-way ANOVA with repeated measures revealed a pronounced hemisphere effect [F(1, 18) = 80, p < 0.0001]. Šídák post-hoc tests further confirmed significant differences in neuron counts between the hemispheres in specific brain regions: PFC (p < 0.0001), BLA (p < 0.001), IC (p < 0.01), PVT (p < 0.001), and the posterior temporal cortex (TEa/ECT/PERI) (p < 0.05). All bar graphs represent mean ± SEM. Asterisks indicate statistically significant p-values (*: p < 0.05, **: p < 0.01, ***: p < 0.001, ****: p < 0.0001). Note: Directional markers indicate the following orientation: A (anterior), P (posterior), L (left), and R (right).

**Extended Figure 6.**
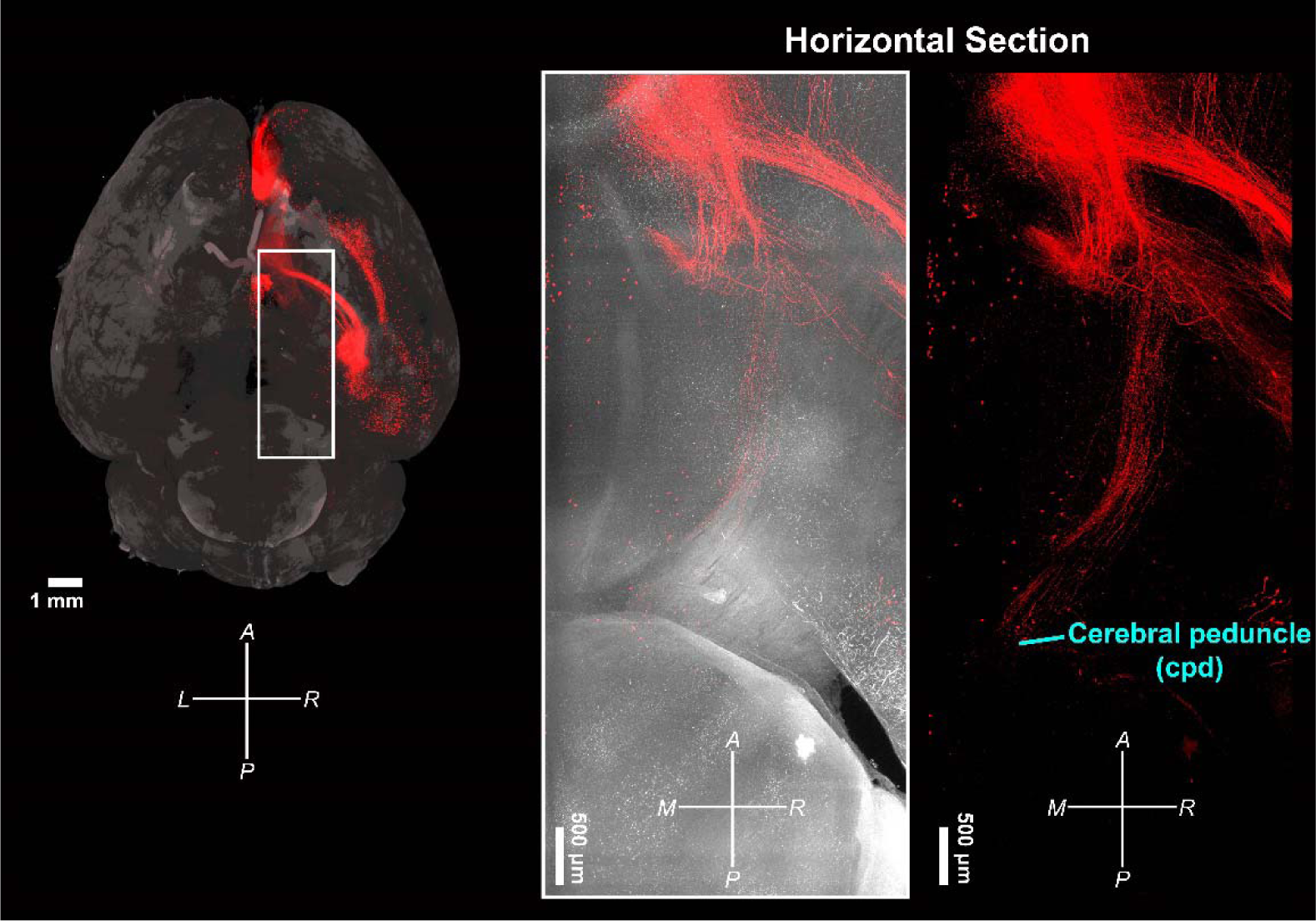
Collateral projection of MNM^PFC→BLA^ to Cerebral Peduncle. (Left) An overview of a cleared brain sample from an Ai-14 mouse with MNM^PFC→BLA^ tracing. The white box highlights the region of interest for the horizontal section. (Middle) A virtual horizontal section displaying the cerebral peduncle (cpd). (Right) A focused view showing only tdTomato labeling in the horizontal section, illustrating the collateral projection of the MNM^PFC→BLA^ to the cpd. Note: Directional markers indicate the following orientation: A (anterior), P (posterior), M (medial), and R (right).

**Extended Figure 7.**
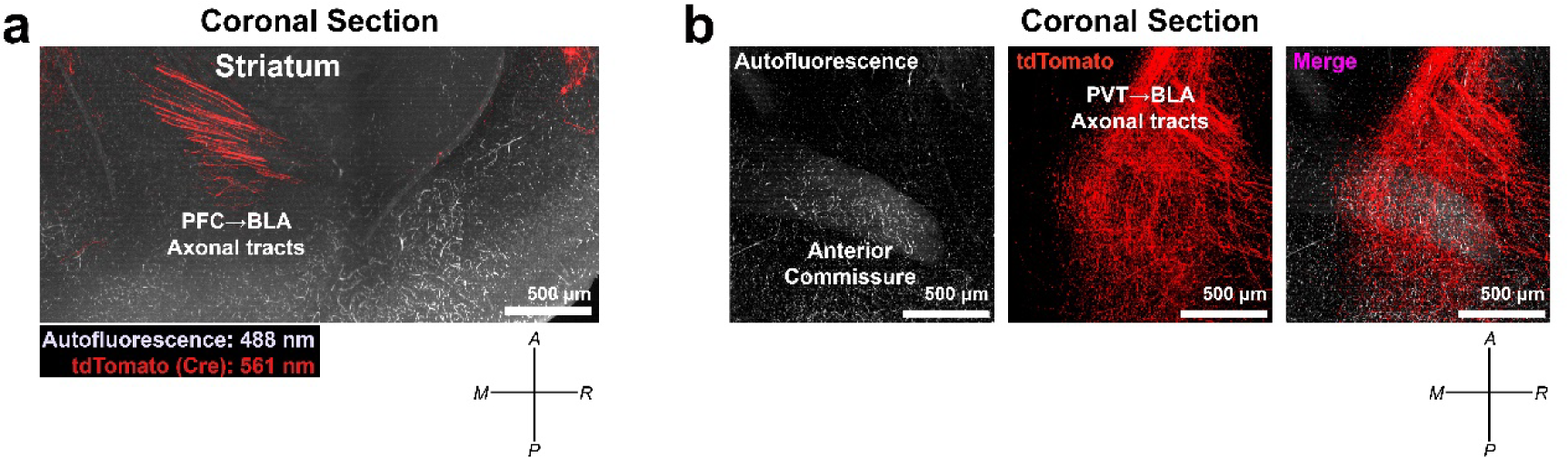
Detailed analysis of axonal connections in MNM^PFC→BLA^. (a) Visualization of PFC→BLA axonal tracts through the striatum. A virtual coronal section from an Ai-14 mouse with MNM^PFC→BLA^ tracing reveals tdTomato-labeled axonal tracts traversing the striatum, illustrating the connection between the PFC and BLA. (b) Complex trajectory of PVT→BLA axonal tracts near the anterior commissure. A virtual coronal section in the region of the anterior commissure showcases the intricate pathway of the tdTomato-labeled PVT→BLA axonal tracts as they navigate around this structure. Note: Directional markers indicate the following orientation: A (anterior), P (posterior), M (medial), and R (right).

**Extended Figure 8.**
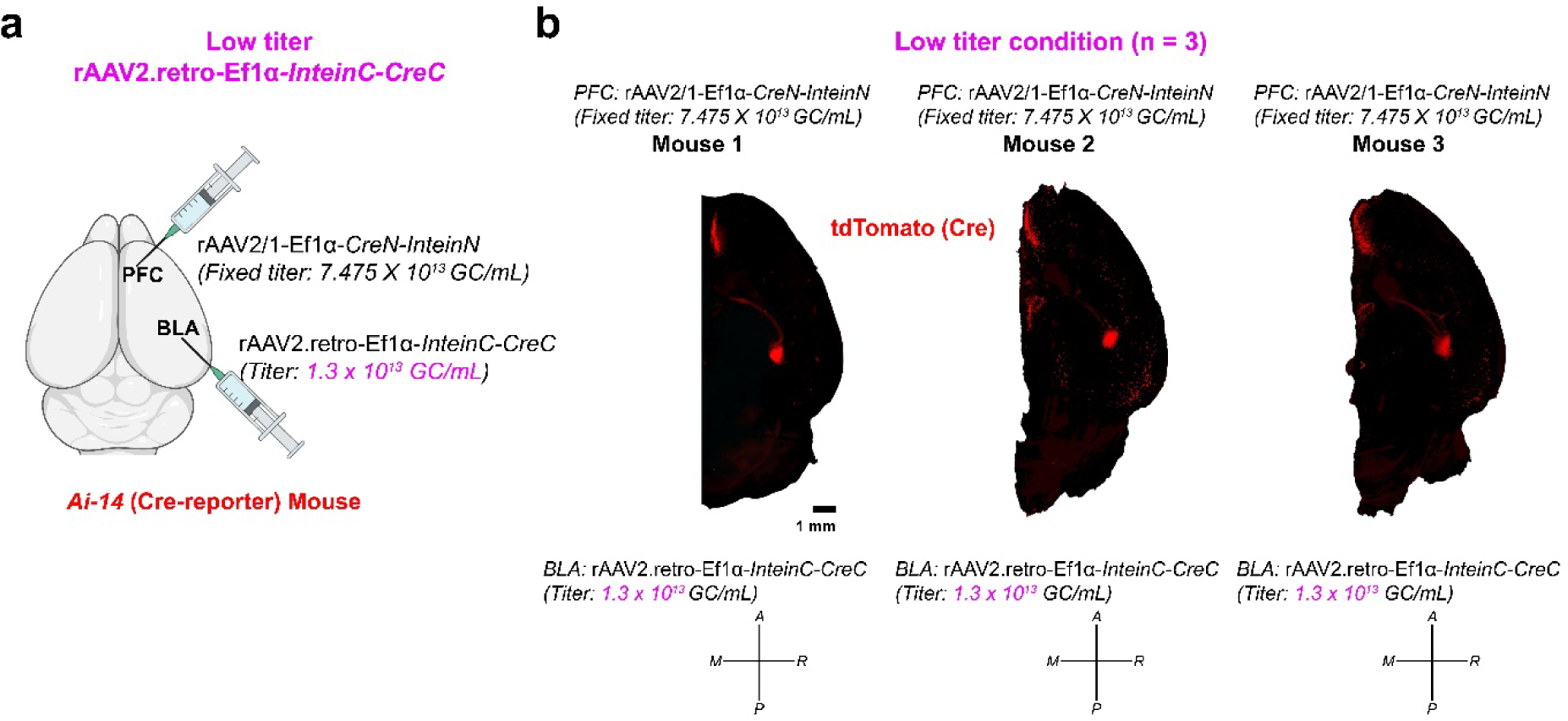
MNM^PFC→BLA^ tracing with reduced rAAV titer. (a) Schematic overview. In this experiment, additional Ai-14 mice (n=3) were used to assess the impact of a lower rAAV2.retro-Ef1α*-InteinC-CreC* titer (1.3 ×10^13^ GC/mL) in the right BLA, while maintaining a constant titer of rAAV2/1-Ef1α*-CreN-InteinN* titer (7.475 ×10^13^ GC/mL) in the right PFC. The objective was to determine how a reduced viral titer affects tracing efficiency. (b) Right-hemisphere of cleared brains. The brain samples exhibited weaker tdTomato signals in comparison to those subjected to higher rAAV titer conditions (Extended Fig. 5), indicating that a reduced rAAV titer results in less efficient labeling of network neurons and affects the clarity of MNM tracing. Note: Directional markers indicate the following orientation: A (anterior), P (posterior), M (medial), and R (right).

**Extended Figure 9.**
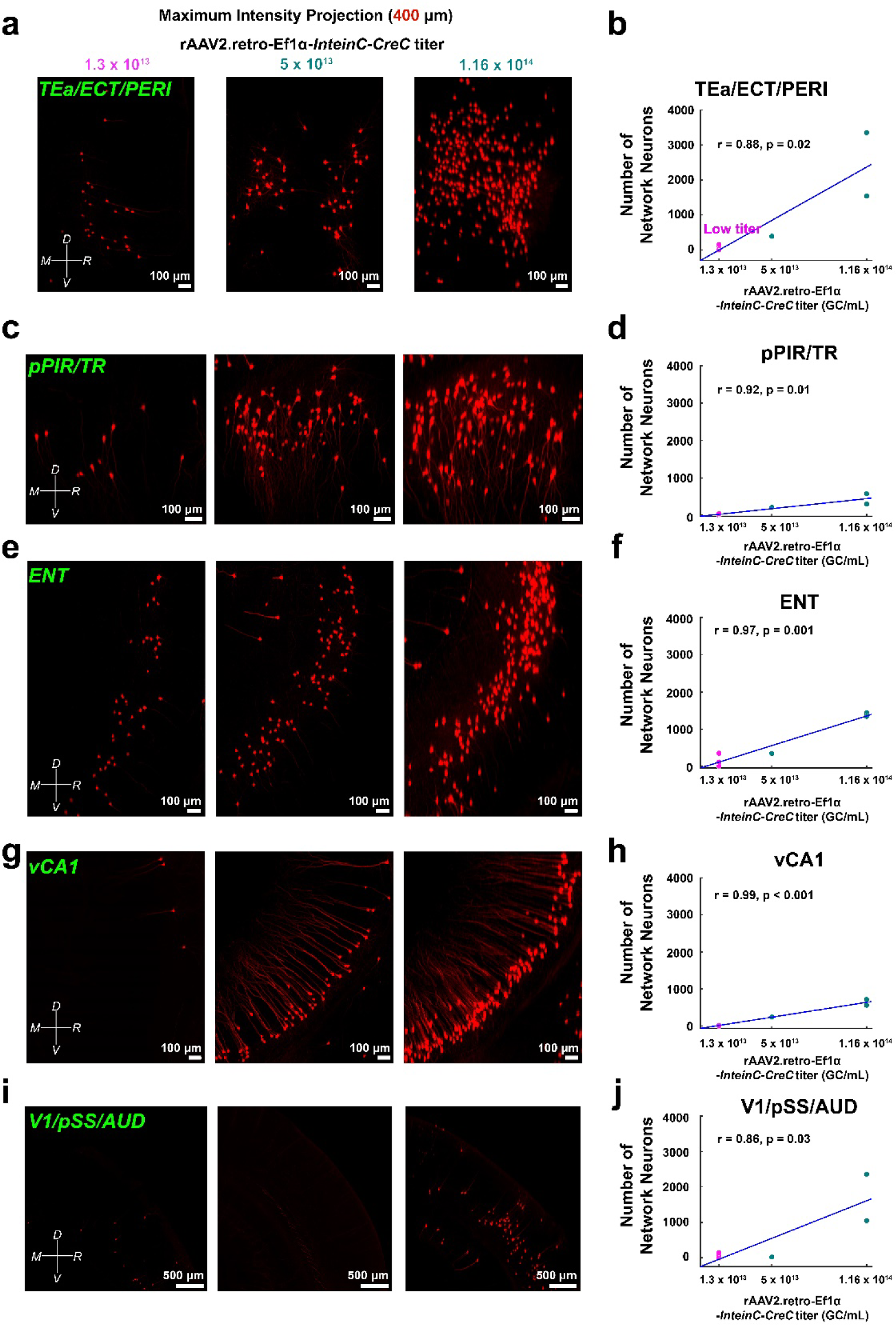
Relationship between rAAV titers and network neuron expression in the posterior hub regions. (a) Visualization of network neurons in the posterior temporal cortex [temporal association areas (TEa)/ ectorhinal area (ECT)/ perirhinal area (PERI)] across varying rAAV2.retro-Ef1α*-InteinC-CreC* titers ranging from 1.3×10^13^ to 5×10^13^ and 1.16×10^14^ GC/mL. Each image is a maximum intensity projection (MIP) from a 400 µm stack. (b) Correlation analysis (Pearson’s) between the posterior temporal cortex neuron expression and rAAV2.retro-Ef1α*-InteinC-CreC* titer, including a least squares error line to depict the positive correlation trend. (c)-(j) Similar visualization and correlation analyses for other posterior hub regions: posterior piriform area (pPIR)/ postpiriform transition area (TR), entorhinal area (ENT), ventral hippocampus CA1 (vCA1), and posterior isocortex [visual area (V1)/ posterior somatosensory areas (pSS)/ auditory areas (AUD)], each with MIPs from 400 µm stacks. Each panel highlights the impact of varying rAAV titers on network neuron expression in these specific areas. Note: Directional markers indicate the following orientation: L (left), R (right), D (dorsal), and V (ventral).

**Extended Figure 10.**
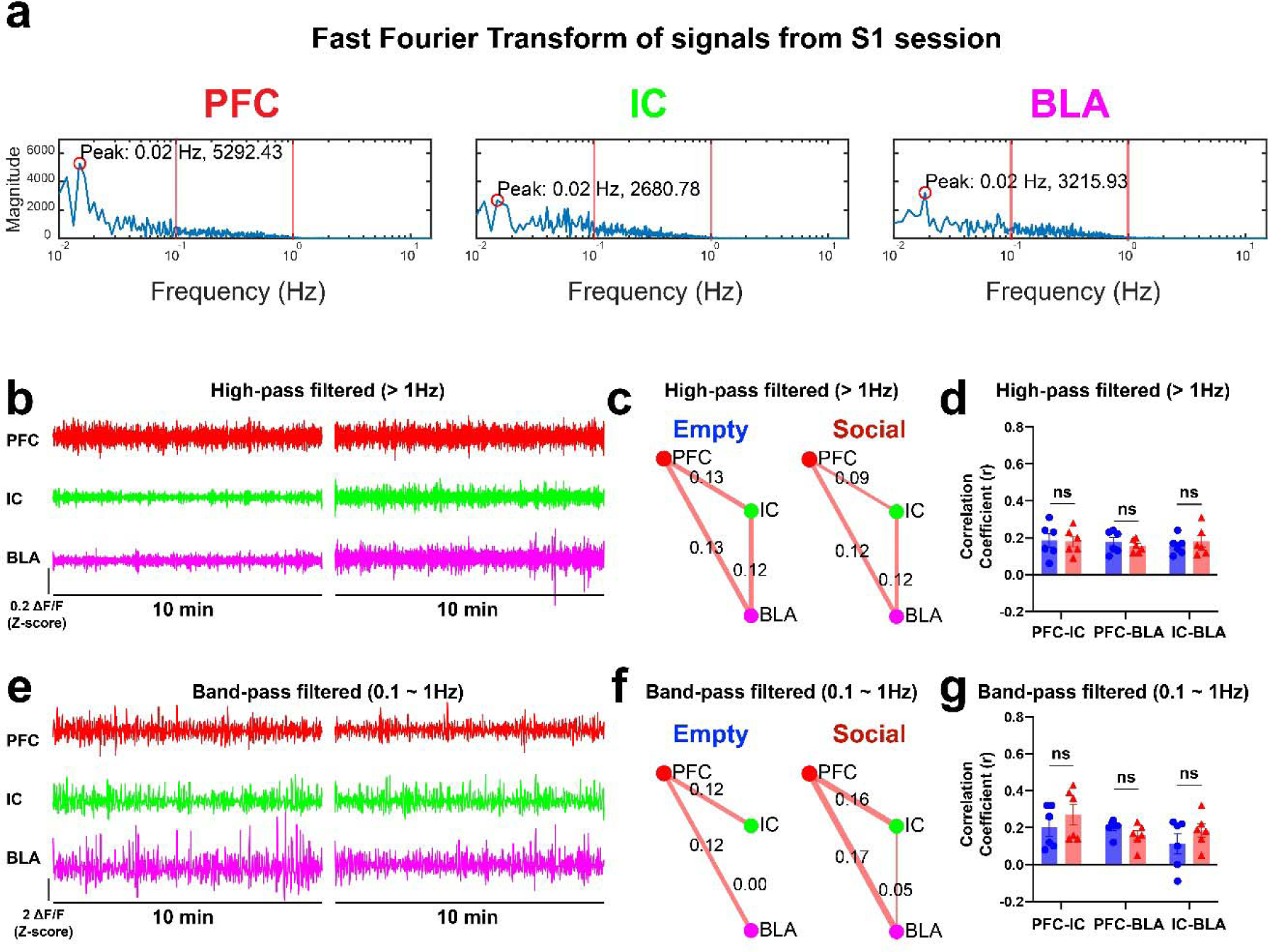
Frequency analysis of fiber photometry signals. (a) Application of Fast Fourier Transform (FFT) on fiber photometry signals from PFC, IC, and BLA. This analysis reveals the highest signal magnitude in the sub-0.1 Hz range, with a decrease in magnitude at higher frequencies. (b) Display of high-pass filtered signals focusing on frequencies above 1 Hz. (c) Correlation analysis of signals above 1 Hz frequency during representative ‘Empty’ and ‘Social’ sessions. (d) Statistical comparison of network correlations for signals above 1 Hz across all ‘Empty’ and ‘Social’ sessions. No significant differences were observed [two-way ANOVA with repeated measures; F(1, 10) = 0.001885, p = 0.97; Šídák post-hoc tests, p > 0.46]. (e-g) Analysis similar to (b-d) but applying band-pass filtering (0.1 ∼ 1 Hz) to the fiber photometry signals. This analysis revealed no significant differences in network correlations at this frequency range [two-way ANOVA with repeated measures; F(1, 10) = 0.6243, p = 0.45; Šídák post-hoc tests, p > 0.20].

## Supplementary Information

**Video 1. Anterograde tracing.**

Anterograde tracing from the right PFC and trans-synaptic tagging of recipient neurons using rAAV2/1-hSyn-Cre

**Video 2. Retrograde tracing**

Retrograde tracing of afferent neurons to the right BLA using rAAV2.retro-hSyn-Cre

**Video 3. Single circuit tracing**

Long-range single circuit tracing from the right PFC to the right BLA

**Video 4. Monosynaptically-interconnected Network Module (MNM) tracing**

Tracing of monosynaptically-interconnected Network Module (MNM) originating from the PFC→BLA connection

**Video 5. Round Social Arena (RSA) test**

Round Social Arena test and simultaneous recording of fiber photometry

**Table 5. Comprehensive details of statistical analyses**

