## Supplementary Table 1 for "Monosynaptically-interconnected Network Module (MNM) Approach for High-Resolution Brain Sub-Network Analysis"

| Figure 3c | Pearson’s Correlation  AAV2.retro titer  X PFC network neurons  (Right hemisphere) | r = 0.9501  95% confidence interval: 0.6054 to 0.9947  R^2^ = 0.9028  P (two-tailed) = 0.0037  P value summary = **  Significant? (alpha = 0.05) = Yes |  |
| --- | --- | --- | --- |
| Figure 3e | Pearson’s Correlation  AAV2.retro titer  X BLA network neurons  (Right hemisphere) | r = 0.9297  95% confidence interval: 0.4813 to 0.9925  R^2^ = 0.8644  P (two-tailed) = 0.0072  P value summary = **  Significant? (alpha = 0.05) = Yes |  |
| Figure 3g | Pearson’s Correlation  AAV2.retro titer  X PVT network neurons  (Right hemisphere) | r = 0.9775  95% confidence interval: 0.8027 to 0.9976  R^2^ = 0.9555  P (two-tailed) = 0.0008  P value summary = ***  Significant? (alpha = 0.05) = Yes |  |
| Figure 3i | Pearson’s Correlation  AAV2.retro titer  X IC network neurons  (Right hemisphere) | r = 0.9525  95% confidence interval: 0.6211 to 0.9950  R^2^ = 0.9073  P (two-tailed) = 0.0033  P value summary = **  Significant? (alpha = 0.05) = Yes |  |
| Figure 4g | Two-way ANOVA with repeated measures  Factor 1: Social stimulus (Empty vs. Social)  Factor 2: Pairing of two brain regions (PFC-IC, PFC-BLA, IC-BLA) | Factor 1: F(1, 15) = 17.34  P = 0.0008  P value summary = ***  Significant? (alpha = 0.05) = Yes  Mean difference (Empty - Social) = -0.1383  95% confidence interval:  -0.2091 to -0.06752  Factor 2: F(2, 15) = 2.869  P = 0.0881  P value summary = ns  Significant? (alpha = 0.05) = No  Factor 1 x Factor 2 interaction:  F(2,15) = 2.507  P = 0.1150  P value summary = ns  Significant? (alpha = 0.05) = No | Post-hoc test (Šidák’s correction)  PFC-IC  Empty vs. Social  Mean difference = -0.2333  95% confidence interval:  -0.3560 to -0.1107  Adjusted P Value = 0.0010  Summary = **  Significant? = Yes  PFC-BLA  Empty vs. Social  Mean difference = -0.05167  95% confidence interval:  -0.3560 to -0.1107  Adjusted P Value = 0.3834  Summary = ns  Significant? = No  IC-BLA  Empty vs. Social  Mean difference = -0.1300  95% confidence interval:  -0.2527 to -0.007348  Adjusted P Value = 0.0392  Summary = *  Significant? = Yes  Empty  PFC-IC vs. PFC-BLA  Mean difference = 0.02000  95% confidence interval:  -0.1431 to 0.1831  Adjusted P Value = 0.9859  Summary = ns  Significant? = No  Empty  PFC-IC vs. IC-BLA  Mean difference = 0.04333  95% confidence interval:  -0.1198 to 0.2064  Adjusted P Value = 0.8801  Summary = ns  Significant? = No  Empty  PFC-BLA vs. IC-BLA  Mean difference = 0.02333  95% confidence interval:  -0.1398 to 0.1864  Adjusted P Value = 0.9781  Summary = ns  Significant? = No  Social  PFC-IC vs. PFC-BLA  Mean difference = 0.2017  95% confidence interval:  0.03857 to 0.3648  Adjusted P Value = 0.0117  Summary = *  Significant? = Yes  Social  PFC-IC vs. IC-BLA  Mean difference = 0.1467  95% confidence interval:  -0.01643 to 0.3098  Adjusted P Value = 0.0882  Summary = ns  Significant? = No  Social  PFC-BLA vs. IC-BLA  Mean difference = -0.05500  95% confidence interval:  -0.2181 to 0.1081  Adjusted P Value = 0.7847  Summary = ns  Significant? = No |
| Figure 4j | Two-way ANOVA with repeated measures  Factor 1: Social stimulus (Empty vs. Social)  Factor 2: Pairing of two brain regions (PFC-IC, PFC-BLA, IC-BLA), < 0.1 Hz | Factor 1: F(1, 15) = 25.42  P = 0.0001  P value summary = ***  Significant? (alpha = 0.05) = Yes  Mean difference (Empty - Social) = -0.2139  95% confidence interval:  -0.3043 to -0.1235  Factor 2: F(2, 15) = 2.725  P = 0.0978  P value summary = ns  Significant? (alpha = 0.05) = No  Factor 1 x Factor 2 interaction:  F(2,15) = 3.721  P = 0.0487  P value summary = *  Significant? (alpha = 0.05) = Yes | Post-hoc test (Šidák’s correction)  PFC-IC  Empty vs. Social  Mean difference = -0.3750  95% confidence interval:  -0.5316 to -0.2184  Adjusted P Value = 0.0001  Summary = ***  Significant? = Yes  PFC-BLA  Empty vs. Social  Mean difference = -0.1083  95% confidence interval:  -0.2649 to 0.04828  Adjusted P Value = 0.1611  Summary = ns  Significant? = No  IC-BLA  Empty vs. Social  Mean difference = -0.1583  95% confidence interval:  -0.3149 to -0.001718  Adjusted P Value = 0.0478  Summary = *  Significant? = Yes  Empty  PFC-IC vs. PFC-BLA  Mean difference = 0.02167  95% confidence interval:  -0.2028 to 0.1594  Adjusted P Value = 0.9869  Summary = ns  Significant? = No  Empty  PFC-IC vs. IC-BLA  Mean difference = -0.02833  95% confidence interval:  -0.2094 to 0.1528  Adjusted P Value = 0.9717  Summary = ns  Significant? = No  Empty  PFC-BLA vs. IC-BLA  Mean difference = -0.006667  95% confidence interval:  -0.1878 to 0.1744  Adjusted P Value = 0.9996  Summary = ns  Significant? = No  Social  PFC-IC vs. PFC-BLA  Mean difference = 0.2450  95% confidence interval:  0.06390 to 0.4261  Adjusted P Value = 0.0055  Summary = **  Significant? = Yes  Social  PFC-IC vs. IC-BLA  Mean difference = 0.1883  95% confidence interval:  0.007231 to 0.3694  Adjusted P Value = 0.0395  Summary = *  Significant? = Yes  Social  PFC-BLA vs. IC-BLA  Mean difference = -0.05667  95% confidence interval:  -0.2378 to 0.1244  Adjusted P Value = 0.8197  Summary = ns  Significant? = No |
| Extended Figure 3c | Paired t test  Number of circuit neuron cell bodies in the PFC  (Left vs. Right hemisphere) | T(2) = 4.6  Mean difference = 4615  95% confidence interval: 298.1 to 8932  R^2^ = 0.9136  P (two-tailed) = 0.0033  P value summary = **  Significant? (alpha = 0.05) = Yes |  |
| Extended Figure 3e | Paired t test  Number of circuit neuron cell bodies in the BLA  (Left vs. Right hemisphere) | T(2) = 3.671  Mean difference = 27.33  95% confidence interval: -4.705 to 59.37  R^2^ = 0.8708  P (two-tailed) = 0.0669  P value summary = ns  Significant? (alpha = 0.05) = No |  |
| Extended Figure 3f | Paired t test  Axonal intensity in the BLA  (Left vs. Right hemisphere) | T(2) = 6.084  Mean difference = 29.01  95% confidence interval: 8.494 to 49.52  R^2^ = 0.9487  P (two-tailed) = 0.0260  P value summary = *  Significant? (alpha = 0.05) = Yes |  |
| Extended Figure 5b | Two-way ANOVA with repeated measures  Factor 1: Hemisphere [Left (contralateral) vs. Right (ipsilateral)]  Factor 2: Brain region | Factor 1: F(1, 18) = 80.00  P < 0.0001  P value summary = ****  Significant? (alpha = 0.05) = Yes  Mean difference = -2204  95% confidence interval:  -2721 to -1686  Factor 2: F(8, 18) = 6.747  P = 0.0004  P value summary = ***  Significant? (alpha = 0.05) = Yes  Factor 1 x Factor 2 interaction:  F(8, 18) = 6.258  P = 0.0006  P value summary = ***  Significant? (alpha = 0.05) = Yes | Post-hoc test (Šidák’s correction)  Hemisphere [Left (contralateral) vs. Right (ipsilateral)]  PFC  Mean difference: -6148  95% confidence interval:  -7701 to -4595  Adjusted P Value < 0.0001  P value summary = ****  Significant? (alpha = 0.05) = Yes  BLA  Mean difference: -2910  95% confidence interval:  -4463 to -1357  Adjusted P Value = 0.0010  P value summary = **  Significant? (alpha = 0.05) = Yes  IC  Mean difference: -2469  95% confidence interval:  -4022 to -916.5  Adjusted P Value = 0.0036  P value summary = **  Significant? (alpha = 0.05) = Yes  PVT  Mean difference: -3634  95% confidence interval:  -5187 to -2081  Adjusted P Value = 0.0001  P value summary = ***  Significant? (alpha = 0.05) = Yes  vCA1  Mean difference: -502.0  95% confidence interval:  -2055 to -1051  Adjusted P Value = 0.5057  P value summary = ns  Significant? (alpha = 0.05) = No  TEa/ECT/PERI  Mean difference: -1688  95% confidence interval:  -3241 to -135.5  Adjusted P Value = 0.0347  P value summary = *  Significant? (alpha = 0.05) = Yes  pPIR/TR  Mean difference: -361.0  95% confidence interval:  -1914 to -1192  Adjusted P Value = 0.6312  P value summary = ns  Significant? (alpha = 0.05) = No  ENT  Mean difference: -1038  95% confidence interval:  -2591 to -514.8  Adjusted P Value = 0.1772  P value summary = ns  Significant? (alpha = 0.05) = No  V1/pSS/AUD  Mean difference: -1081  95% confidence interval:  -2634 to -471.5  Adjusted P Value = 0.1607  P value summary = ns  Significant? (alpha = 0.05) = No |
| Extended Figure 9b | Pearson’s Correlation  AAV2.retro titer  X TEa/ECT/PERI network neurons  (Right hemisphere) | r = 0.8834  95% confidence interval: 0.2539 to 0.9872  R^2^ = 0.7804  P (two-tailed) = 0.0196  P value summary = *  Significant? (alpha = 0.05) = Yes |  |
| Extended Figure 9d | Pearson’s Correlation  AAV2.retro titer  X pPIR/TR network neurons  (Right hemisphere) | r = 0.9160  95% confidence interval: 0.4068 to 0.9909  R^2^ = 0.8390  P (two-tailed) = 0.0103  P value summary = *  Significant? (alpha = 0.05) = Yes |  |
| Extended Figure 9f | Pearson’s Correlation  AAV2.retro titer  X ENT network neurons  (Right hemisphere) | r = 0.9692  95% confidence interval: 0.7383 to 0.9967  R^2^ = 0.9393  P (two-tailed) = 0.0014  P value summary = **  Significant? (alpha = 0.05) = Yes |  |
| Extended Figure 9h | Pearson’s Correlation  AAV2.retro titer  X vCA1 network neurons  (Right hemisphere) | r = 0.9871  95% confidence interval: 0.8824 to 0.9986  R^2^ = 0.9743  P (two-tailed) = 0.0002  P value summary = ***  Significant? (alpha = 0.05) = Yes |  |
| Extended Figure 9j | Pearson’s Correlation  AAV2.retro titer  X V1/pSS/AUD network neurons  (Right hemisphere) | r = 0.8557  95% confidence interval: 0.1409 to 0.9838  R^2^ = 0.7306  P (two-tailed) = 0.0301  P value summary = *  Significant? (alpha = 0.05) = Yes |  |
| Extended Figure 10d | Two-way ANOVA with repeated measures  Factor 1: Social stimulus (Empty vs. Social)  Factor 2: Pairing of two brain regions (PFC-IC, PFC-BLA, IC-BLA), > 1 Hz | Factor 1: F(1, 15) = 0.005540  P = 0.9416  P value summary = ns  Significant? (alpha = 0.05) = No  Mean difference (Empty - Social) = 0.001111  95% confidence interval:  -0.03071 to 0.03293  Factor 2: F(2, 15) = 0.1441  P = 0.8669  P value summary = ns  Significant? (alpha = 0.05) = No  Factor 1 x Factor 2 interaction:  F(2,15) = 0.7742  P = 0.4786  P value summary = ns  Significant? (alpha = 0.05) = No | Post-hoc test (Šidák’s correction)  PFC-IC  Empty vs. Social  Mean difference = 0.005000  95% confidence interval:  -0.05011 to 0.06011  Adjusted P Value = 0.8493  Summary = ns  Significant? = No  PFC-BLA  Empty vs. Social  Mean difference = 0.02167  95% confidence interval:  -0.03344 to 0.07678  Adjusted P Value = 0.4152  Summary = ns  Significant? = No  IC-BLA  Empty vs. Social  Mean difference = -0.02333  95% confidence interval:  -0.07844 to 0.03178  Adjusted P Value = 0.3811  Summary = ns  Significant? = No  Empty  PFC-IC vs. PFC-BLA  Mean difference = 0.008333  95% confidence interval:  -0.08571 to 0.1024  Adjusted P Value = 0.9741  Summary = ns  Significant? = No  Empty  PFC-IC vs. IC-BLA  Mean difference = 0.02833  95% confidence interval:  -0.06571 to 0.1224  Adjusted P Value = 0.7403  Summary = ns  Significant? = No  Empty  PFC-BLA vs. IC-BLA  Mean difference = 0.02000  95% confidence interval:  -0.07404 to 0.1140  Adjusted P Value = 0.8601  Summary = ns  Significant? = No  Social  PFC-IC vs. PFC-BLA  Mean difference = 0.025000  95% confidence interval:  -0.06904 to 0.1190  Adjusted P Value = 0.7908  Summary = ns  Significant? = No  Social  PFC-IC vs. IC-BLA  Mean difference = 0.000  95% confidence interval:  -0.09494 to 0.09494  Adjusted P Value > 0.9999  Summary = ns  Significant? = No  Social  PFC-BLA vs. IC-BLA  Mean difference = -0.025000  95% confidence interval:  -0.1190 to 0.06904  Adjusted P Value = 0.7908  Summary = ns  Significant? = No |
| Extended Figure 10g | Two-way ANOVA with repeated measures  Factor 1: Social stimulus (Empty vs. Social)  Factor 2: Pairing of two brain regions (PFC-IC, PFC-BLA, IC-BLA), 0.1 ~ 1 Hz | Factor 1: F(1, 15) = 1.643  P = 0.2193  P value summary = ns  Significant? (alpha = 0.05) = No  Mean difference (Empty - Social) =  -0.03333  95% confidence interval:  -0.08876 to 0.02209  Factor 2: F(2, 15) = 1.591  P = 0.2362  P value summary = ns  Significant? (alpha = 0.05) = No  Factor 1 x Factor 2 interaction:  F(2,15) = 2.080  P = 0.1595  P value summary = ns  Significant? (alpha = 0.05) = No | Post-hoc test (Šidák’s correction)  PFC-IC  Empty vs. Social  Mean difference = -0.07000  95% confidence interval:  -0.1660 to 0.02600  Adjusted P Value = 0.1410  Summary = ns  Significant? = No  PFC-BLA  Empty vs. Social  Mean difference = 0.04167  95% confidence interval:  -0.05433 to 0.1377  Adjusted P Value = 0.3695  Summary = ns  Significant? = No  IC-BLA  Empty vs. Social  Mean difference = -0.07167  95% confidence interval:  -0.1677 to 0.02433  Adjusted P Value = 0.1324  Summary = ns  Significant? = No  Empty  PFC-IC vs. PFC-BLA  Mean difference = 0.000  95% confidence interval:  -0.1492 to 0.1492  Adjusted P Value > 0.9999  Summary = ns  Significant? = No  Empty  PFC-IC vs. IC-BLA  Mean difference = 0.08833  95% confidence interval:  -0.06085 to 0.2375  Adjusted P Value = 0.3745  Summary = ns  Significant? = No  Empty  PFC-BLA vs. IC-BLA  Mean difference = 0.08833  95% confidence interval:  -0.06085 to 0.2375  Adjusted P Value = 0.3745  Summary = ns  Significant? = No  Social  PFC-IC vs. PFC-BLA  Mean difference = 0.1177  95% confidence interval:  -0.03752 to 0.2608  Adjusted P Value = 0.1907  Summary = ns  Significant? = No  Social  PFC-IC vs. IC-BLA  Mean difference = 0.08667  95% confidence interval:  -0.06252 to 0.2358  Adjusted P Value = 0.3908  Summary = ns  Significant? = No  Social  PFC-BLA vs. IC-BLA  Mean difference = -0.02500  95% confidence interval:  -0.1742 to 0.1242  Adjusted P Value = 0.9656  Summary = ns  Significant? = No |
